# Encoding of bimanual movement directions across the human sensorimotor system

**DOI:** 10.64898/2026.09.18.752753

**Authors:** Ali Ghavampour, Atsushi Yokoi, Diogo F. Duarte, Jean-Jacques Orban de Xivry, J. Andrew Pruszynski, Jörn Diedrichsen

## Abstract

Bimanual coordination requires the control of each hand to account for movements of the other hand. To enable such coordination, it has been hypothesized that the brain has a representation of bimanual movements that supersedes a linear combination of the unimanual movements. To test for such a representation, we mapped the encoding of movement direction during unimanual and bimanual wrist movements in the human brain using functional magnetic resonance imaging (fMRI). For unimanual movements, we found that both the contralateral and ipsilateral movement directions were represented across most cortical motor regions. The representations were similar for mirror-symmetric movements across the hands, revealing an intrinsic (body-centric) code in premotor and parietal regions. For bimanual movements, the neural representations of movement directions could be explained by three components. First, the contralateral movement direction was represented as in the unimanual condition, whereas the representation of the ipsilateral movement direction disappeared. Second, in higher-order regions (rostral premotor and posterior parietal cortex) we found relatively elevated activity when the two movements were directed in unrelated directions. This activity was likely caused by the extra effort to deal with incongruent spatial targets across the hands. Third, and independent of this congruency effect, we found a non-linear interaction between contra- and ipsilateral movement that encoded the specific bimanual movement combination across sensorimotor, premotor, and parietal regions. This representation was suited to facilitate bimanual coordination and learning.

## Introduction

Many daily actions depend on simultaneous movements of both hands. Sometimes the hands need to move independently, such as stirring a pot with one hand and adding the next ingredients with the other hand. This requires the motor system to control each hand without being disturbed by what the other hand is doing. Often, however, we are trying to achieve a single goal with coordinated movements of both hands, like opening a bottle or folding laundry (Swinnen and Wenderoth, 2004), which requires that the neural controller of each hand takes into account the state and movements of the other hand. How is this achieved in the human brain?

Behavioural studies of bimanual coordination have focused on two core observations. First, even though the hands can move independently when trying to achieve different goals, bimanual interference can be observed when these two goals are incongruent. For example, it is more difficult to produce different spatial movements with the two hands than the same or mirror symmetric spatial movements (Franz et al., 1991; Swinnen et al., 1998, 2001, 2002; Albert and Ivry, 2009). Similarly, producing bimanual movements with different speeds, extents, or frequency is much harder than producing matching movements with both hands (Kelso et al., 1979; Kelso, 1984; Marteniuk et al., 1984; Heuer et al., 1998; Mechsner et al., 2001; Blinch et al., 2014). These observations clearly show that the neural controllers for the two hands interact with each other. Second, it can be shown that the two hands can coordinate their movements with each other when they are trying to achieve a single goal together. When one hand lifts a weight from the other’s palm, the postural hand anticipatorily compensates for the impending load change (Diedrichsen et al., 2003b, 2005) – this is absent when the lift is externally generated. Additionally, the adaptation to force fields in one arm depends partly on the movement direction of the other arm (Yokoi et al., 2011, 2014; Omrani et al., 2013; Orschiedt and Franklin, 2023; Desrochers et al., 2026). The observation that a learned forcefield generalizes as a smooth function of the other arm’s movement direction implies that the neural controller supporting each arm cannot be tuned to either arm alone, rather must also depend on the combination of the two movements. Together, these findings suggest that the brain maintains not only independent representations of each unimanual movement, but also a representation of the specific bimanual combination of movements.

Even though the human sensorimotor cortex mainly controls the contralateral hand (Penfield and Boldrey, 1937; Soteropoulos et al., 2011), neural studies have shown that it represents movements of either hand. Monkey electrophysiology and imaging studies find activity and encoding of ipsilateral hand movement (Cisek et al., 2003; Ganguly et al., 2009; Haar et al., 2017; Ames and Churchland, 2019; Berlot et al., 2019; Heming et al., 2019; Cross et al., 2020; Willett et al., 2020; Guan et al., 2022). Importantly, bimanual movement representation seems not to be a simple additive combination of contralateral and ipsilateral representations. Single cell electrophysiology during bimanual reaching movements in the monkey shows that neurons respond during bimanual movements in a way that is not predicted by the sum of their responses to the constituent unimanual movements (Donchin et al., 1998, 2001, 2002; Steinberg et al., 2002; Rokni et al., 2003). In humans, an fMRI study of bimanual finger movements found similar non-additive encoding, with dorsal premotor cortex representing the specific bimanual combination rather than linear combination of the constituent unimanual movements (Diedrichsen et al., 2013).

A similar study for bimanual goal-directed reaching movements in humans is currently missing. Despite the evidence for contra- and ipsilateral representations of movements, it is still unknown how 1) unimanual representations of goal-directed reaching movements interact during bimanual movements; and 2) how these representations distribute in multiple cortical regions. Here we designed an fMRI-compatible spatial bimanual task that required independent movements of the left and right wrist to six spatial targets. For unimanual movements, we were able to map the encoding of both the contralateral and ipsilateral movement direction across cortical motor areas. By measuring the activity patterns of all 36 combinations of left and right targets, we were then able to distinguish different models of how bimanual movements are encoded in these regions. Specifically, we were able to dissociate activity changes that are caused by the requirement to move the two hands in different directions (Sadato et al., 1997; Debaere et al., 2004; Wenderoth et al., 2004, 2005) from genuine encoding of the bimanual movement combination, which would be required for detailed bimanual coordination.

## Methods

### Participants

Twelve healthy self-reported right-handed participants (mean age = 26.7 years, SD = 4.2, range = 18-35; 5 females) were recruited. All procedures were designed in accordance with the Declaration of Helsinki and approved by the Western University Research Ethics Board, and the Research Ethics Board of University College London.

### Apparatus

The wrist device featured a fencing grip mounted to allow rotation along two axes: vertical rotation (wrist ulnar and radial deviation) and horizontal rotation (wrist flexion and extension) (Fig. 1A). The axes were slightly offset from each other to match the natural axes of rotation in the wrist (Crisco et al., 2011). The handles were held with the thumb pointing upwards. Each axis was continuously recorded by a Bourns 3382H potentiometer at a sampling rate of 200 Hz. Horizontal and vertical wrist rotations were mapped to position of a cursor along the x- and y-axes on the screen, respectively. Flexion and extension of the wrist led to horizontal movements of the cursor, and pronation and supination to vertical movement. Home position of the cursor was calibrated to each participant’s resting wrist posture, such that any deviation from this position displaced the cursor from center, and returning to the resting posture re-centered the cursor.

**Figure 1:**
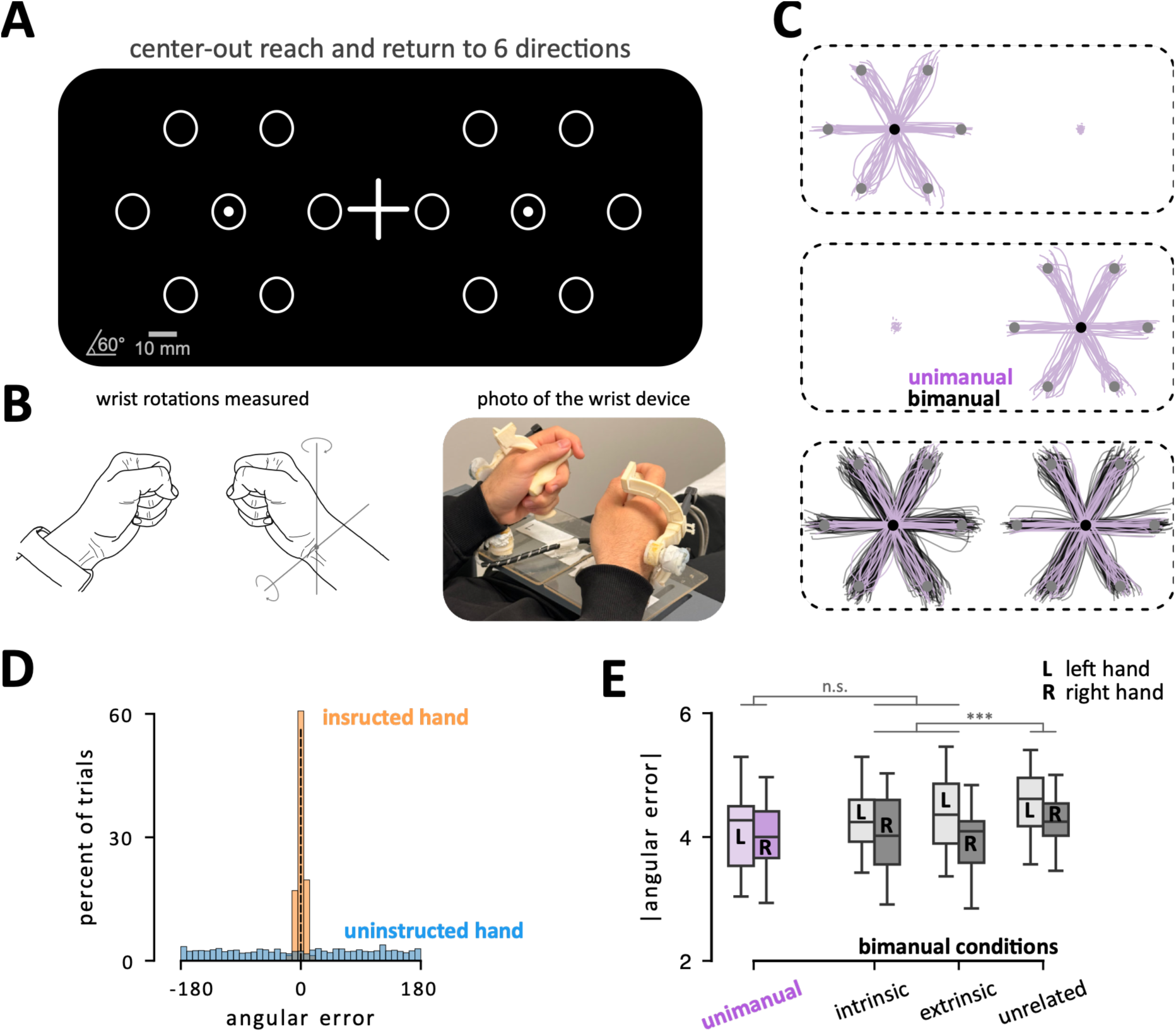
Task & performance. **A)** What participants see on the screen at all times (possible target locations at 0, 60, etc.). The small circles are hand cursors and “+” is the fixation cross. The measurements in the bottom left corner are solely for visualization in this paper. **B)** The drawing of the hands shows the neutral position and the axes of rotations. A photograph of the device held in the neutral position is provided. **C)** Reach trajectories from a representative participant in the scanner. Unimanual trajectories (pink) are overlaid on bimanual trajectories (black) for the same hand and the same targets. **D)** Angular error at peak reach amplitude for the instructed (orange) and uninstructed (blue) hand during unimanual conditions. **E)** Absolute angular error for the left (L) and right (R) hand across unimanual and three types of bimanual conditions (intrinsic, extrinsic, incongruent).

The screen displayed the cursors, the fixation cross, the two home positions, and six potential target locations for each hand at all times (Fig. 1A). The fixation cross (20 mm width and height) was displayed in the center of the screen. The two home positions for the left and the right hand (circles of 5 mm radius) were presented 55 mm to the left and right of a fixation cross. The target locations were unfilled circles with a radius of 5 mm, radially arranged at 0°, 60°, 120°, 180°, 240°, and 300° at 35 mm (center-to-center) from the home position. During the fMRI experiment, visual stimuli were rear-projected onto a screen and viewed via a mirror mounted on the head coil, such that 10 mm on the screen corresponded to approximately 0.87 degrees of visual angle.

### Task

Participants were instructed to keep their gaze on a central fixation cross throughout each trial. At the start of each trial, participants had to keep both left and right hand cursors in the home positions. At the beginning of the planning phase, one or two targets was filled with white indicating the target(s) and whether a unimanual or bimanual movement was required. During the 2000 ms of preparation, participants were instructed to fixate on the central fixation cross and prepare the upcoming movement. At the beginning of the movement phase, the fixation cross turned green signalling the go-cue. Over the subsequent 4500 ms participants had to leave the home position(s), reach the target(s), and return to the home position all in a single continuous movement. For the home position and targets we introduced an extra tolerance. Participants had to be within a 8.5mm radius of the center of the home position and within a 10.6mm radius of the target center to count as a success. If participants left the home position before the go-cue, left the home position more than 800ms after go-cue, missed the target(s), moved an uninstructed hand out of the home position, or did not return to the home position within 4500 ms, the trial was counted as an error. Any error turned the fixation cross blue for the remainder of the trial without interrupting it. After finishing the trial, participants received feedback on the trial score (maximum 1 point for unimanual and maximum 2 points for bimanual) and total score of the run.

### Experiment design

Participants completed all possible unimanual (6 left-hand + 6 right-hand = 12 conditions) and bimanual (6 × 6 = 36 conditions) movement conditions. Each of these 48 conditions was performed twice in each imaging run. The sequence of the 96 trials in each run was fully randomized for each participant. Each participant underwent two imaging sessions with a total of 10 runs (approximately 4 hours per person). Within each run, trials were presented every 7 seconds, with 9 randomly selected trials extended by an additional 5 seconds of rest to improve baseline BOLD signal estimation. At the end of each run participants remained still in the home position for an additional 10 seconds to capture the slow hemodynamic response function (HRF) related to the last trial. Between runs, participants were allowed to rest for 2-4 minutes. The unimanual and bimanual conditions had a success rate of 96.7% and 83.6%, respectively. To familiarize the participants with the task and setup, we trained each participant in a supine position for 10 runs in a mock scanner before the first scanning session.

### Behavioral analysis

We used the position of the cursor at the go-cue as the start position for that trial. We then determined the moment in which the trajectory maximally deviated from that position. The distance from home target was used as the movement amplitude, and the direction was compared to the target direction to measure the angular error in degrees. This analysis was performed on both hands, whether a movement of that hand was instructed or not.

We measured reaction time (RT) as the time from the go cue until either hand left the home position (3.5 mm away from center). Furthermore, movement time (MT) was defined as the time from RT until the target(s) had been acquired and both hands had returned to their home positions.

### fMRI data acquisition

Data for five of the twelve participants were collected at Western University. We used a 3 Tesla Siemens MAGNETOM Prisma scanner with a 32-channel head coil, at the Centre for Functional and Metabolic Mapping located at Western university. Blood oxygenation level-dependent (BOLD) response was measured while participants were doing the experiment in a supine position. In each run 740 functional images were obtained using a multi-band 2D echoplanar imaging sequence with a TR repetition time of 1000ms. The other seven participants were collected at Wellcome Trust UCL using a Siemens Trio 3 Tesla MR system in a similar setup with TR of 2720ms, resulting in 269 images per run. Per image, we acquired 32 slices in an interleaved sequence at a thickness of 2.3mm, 0% gap (2mm, 15% gap for UCL) and an in-plane resolution of 2.3 x 2.3 mm^2^. To account for the magnetic field inhomogeneities we acquired field map at each scanning session. Furthermore, a T1-weighted anatomical scan was obtained using MPRAGE (magnetization-prepared rapid gradient echo) sequence with a voxel size of 1mm isotropic.

### Preprocessing and first-level analysis

Using SPM12 (https://www.fil.ion.ucl.ac.uk/spm/), functional images were realigned and field map corrected (unwarped) to correct for head motion and magnetic field inhomogeneities. The functional images were then co-registered to each participant’s anatomical image. Next, the minimally preprocessed data were analyzed using a general linear model (GLM; Friston et al., 1994). Per run, each of 48 conditions (6 unimanual left, 6 unimanual right, 36 bimanual) was modeled with a separate regressor. Each event was modelled with a boxcar function beginning at target presentation and lasting 2500 ms. These regressors were convolved with a hemodynamic response function fitted per participant maximizing the fit quality of a model that only has three regressors denoting contralateral movement, ipsilateral movement, or bimanual movement regardless of their directions. An intercept regressor was also added for each run. The first-level GLM therefore estimated the size of the evoked activity for each condition (*β* weights). For visual display and univariate analyses (Fig. 2), we calculated the percent signal change (PSC) for each condition relative to the baseline activation (rest) for each voxel and condition. The PSC values were then averaged across all runs. In the results section the PSC is referred to as *activation* estimates.

**Figure 2:**
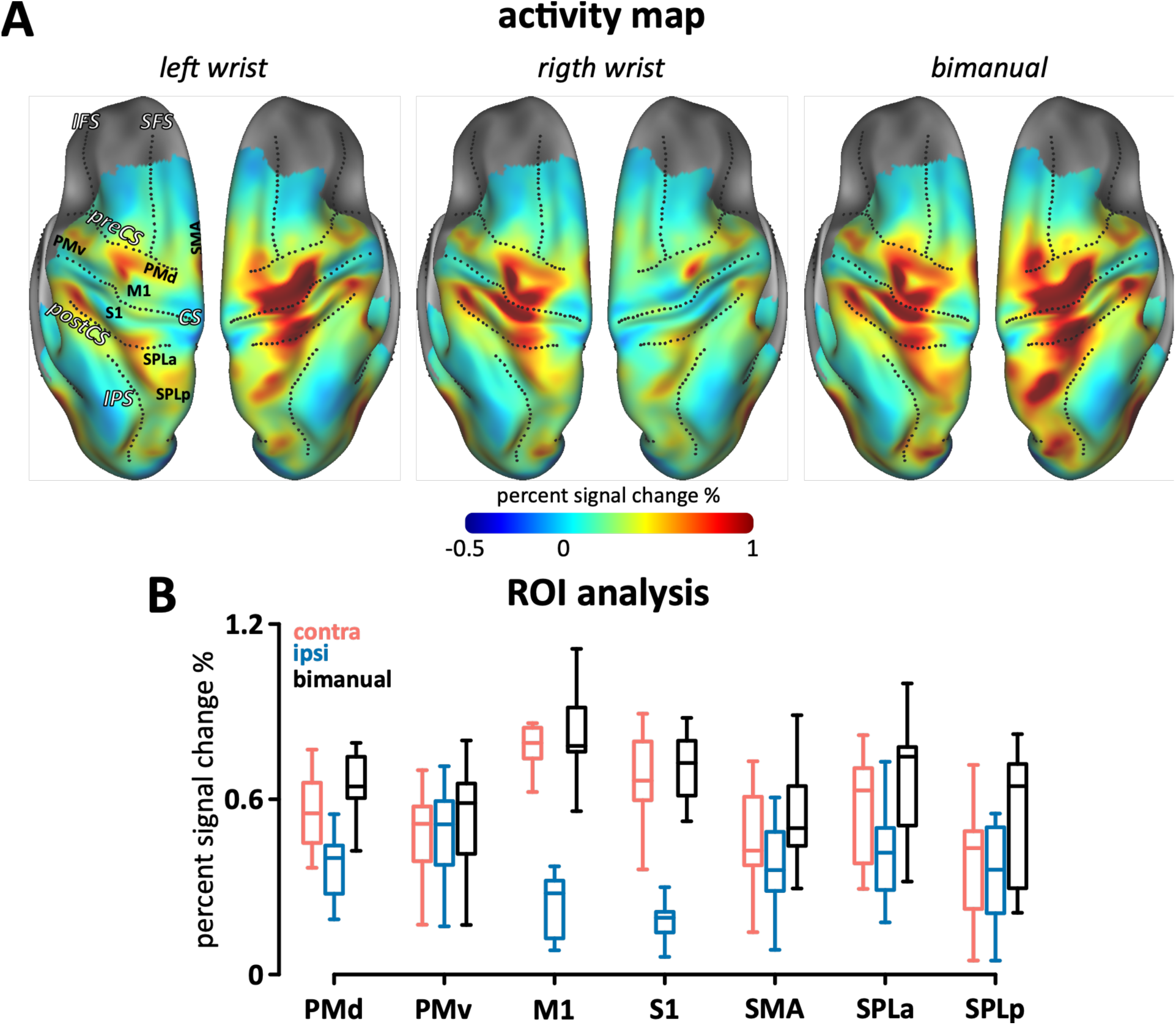
BOLD increase during unimanual conditions. **A)** Group-averaged percent signal change (PSC) mapped across the cortical surface, shown for the left and right hemisphere. Approximate locations of regions of interest (ROIs) are labeled. Anatomical surfaces are symmetric between hemispheres. **B)** Percent signal change within ROIs for contralateral (red), ipsilateral (blue) and bimanual (black) wrist movements.

### Surface-based analysis

We used Freesurfer (Dale et al., 1999; Fischl et al., 1999a) to reconstruct the participant’s pial and white/gray matter surfaces based on their T1-weighted anatomical image. The individual surfaces were then aligned to the Freesurfer average atlas (Fischl et al., 1999b). Finally, surfaces were resampled to a common left-right symmetric template (fs_LR32k; (Van Essen et al., 2012)), which allowed projection of each participant’s volume data onto a group surface map.

Individual PSC and contrast maps were projected from native functional volume space onto the cortical surface using subject-specific white-matter and pial boundary meshes (*surfAnalysisPy*; https://github.com/DiedrichsenLab/surfAnalysisPy). For each surface vertex, volume values were sampled at six equally spaced depths between the white-matter and pial surfaces and then averaged the data across depths. Before group analysis, each participant’s surface map was smoothed with a Gaussian kernel (FWHM = 6 mm) using the Connectome Workbench (*wb_command “cifti-smoothing”*).

### ROI definition

Seven Anatomical Regions of Interest (ROIs) were defined, including primary motor (M1) and somatosensory cortex (S1), ventral and dorsal premotor cortex (PMv and PMd), the anterior and posterior aspects superior parietal lobules (SPLa and SPLp), and the supplementary motor areas (SMA). ROI definition was based on a probabilistic cytoarchitectonic atlas (Fischl et al., 2008) with a procedure established in previous work (Kornysheva et al., 2013; Wiestler and Diedrichsen, 2013; Arbuckle et al., 2019; Berlot et al., 2020; Ariani et al., 2025). We selected the part of M1 and S1 that is related to hand and wrist movements by restricting the cytoarchitectonic ROI to 2 cm above and below the hand knob. Broadman area 6 was subdivided in ventral, dorsal and medial aspect.

For data visualization, we also conducted a continuous version of the ROI analysis. We defined a strip on the anatomical surface that includes the sensorimotor ROIs from the rostral PMd to the caudal part of SPLp (Fig. 7). For each participant and hemisphere, we projected the surface data onto a line that ran anterior to posterior through these ROIs. Data was then divided into 3 mm bins. Group profile was then computed by first averaging across the left and right hemispheres within each participant and then averaging across participants.

### Searchlight analysis

Additional to the ROI analyses, we performed a continuous searchlight analysis (Oosterhof et al., 2011). We defined a searchlight as a circle around each surface node using the Dijsktra distance on the individual surface. We then assigned all voxels that lay between the white-gray matter and pial surface to the searchlight. We increased the radius of each searchlight until each searchlight included exactly 200 voxels. This resulted in an average radius of 13.7 mm for the Western and 15.9 mm for the UCL data. The subsequent multivariate analyses were performed on each of the 200 voxels. The result was then assigned to the center node. As the univariate contrast maps, each participant’s map was smoothed with a Gaussian kernel (FWHM = 6 mm) before averaging them to obtain a group map.

To perform the searchlight analysis, we used the Python *AnatSearchlight* toolbox by the Diedrichsen Lab (https://github.com/DiedrichsenLab/AnatSearchlight), and for smoothing and visualization we used *Connectome Workbench v2.1.0, wb_command* and *wb_view* tools by the Human Connectome Project (Marcus et al., 2011).

### Multivariate fMRI analysis and encoding

For all the multivariate analyses, the *β* weights of each voxel were first divided by the root mean square error from the first-level GLM to univariately prewhiten the signal (Walther et al., 2016). We used cross-validated Euclidean distance (Diedrichsen et al., 2020) on these activity estimates to obtain an unbiased estimate of the distance between the fMRI activity patterns for different conditions. If distances are calculated without cross-validation, they always will be positive because the two average activity patterns will differ due to measurement noise, even if the true underlying activity patterns are perfectly identical. To account for this, we estimated the squared distance of condition *i* and *j* (*d*^2^) as follows:

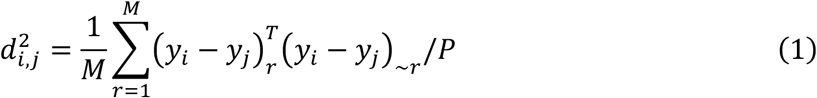

Where M is the number of runs, *y_i_* is the activity pattern of condition *i*, a P-dimensional vector, with *P* being the number of voxels. Here, the activity patterns from run *r* are multiplied to the average of all runs except *r* (∼*r*), such that the noise within a run does not bias the distance estimation. The estimate is averaged across all the leave-one-out folds, resulting in a cross-validated distance estimate of conditions *i* and *j*. This also allows for the distance estimate to be negative. If the two patterns are identical, the expected value of the distance is 0, independent of the amount of noise in the data (Diedrichsen et al., 2020).

Within each participant, we calculated the unbiased distance between every pair of conditions, resulting in a 48 by 48 distance matrix for each ROI (or searchlight). To assess the strength of unimanual and bimanual encoding, we averaged the distance estimates of the 15 (6 x 5 / 2) unimanual contralateral and 15 ipsilateral conditions as well as 630 (36 x 35 / 2) bimanual ones. We used the *Pattern Component Modelling* toolbox (Diedrichsen et al., 2018; https://github.com/DiedrichsenLab/PcmPy) to perform this analysis. The distances were then square root transformed maintaining the sign 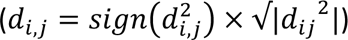 for statistical testing and visual display.

### Correlation between contralateral and ipsilateral activity patterns

To determine the correlation between the unimanual patterns associated with contralateral and ipsilateral movement, we first computed the average activity patterns: Within each participant and ROI (or searchlight), fMRI activity patterns for each of the six movement directions were averaged across runs separately for the contralateral and ipsilateral hand, yielding two 6 × P-voxels matrices. The across-condition mean was removed from each voxel, and the centered matrices were vectorized, and Pearson correlation was computed.

This was done twice with different condition orderings. For the *intrinsic correlation*, contra- and ipsilateral conditions were matched by the movement of the wrist (e.g., left-wrist flexion paired with right-wrist flexion). For the *extrinsic correlation*, they were matched simply by reaching direction in external space (e.g., left-wrist reaching to 60° paired with right-wrist reaching to 60°).

### Noise-corrected correlation of marginal and unimanual contralateral activity patterns

To determine the correlation between the bimanual and unimanual patterns associated with the contralateral movement, we first calculated the marginal bimanual activity patterns. For each run and movement direction of the contralateral hand, we averaged the bimanual activity patterns across the 6 possible movement direction of the ipsilateral hand. Within each ROI, we then compared these 6 marginal bimanual activity patterns with the unimanual patterns for same contralateral movement direction.

Our aim was to assess whether the marginal and unimanual contralateral activity patterns are identical. Estimating the simple Pearson correlation between the patterns, would underestimate the true correlation due to the noisiness of the data (Spearman, 1904). To correct for that, we can derive the maximum-likelihood estimate (MLE) for the correlation with the assumption that both the signal and the measurement noise are Gaussian (Diedrichsen et al., 2026).

To test whether these noise-corrected correlation are significantly smaller than a near-perfect correlation (e.g., *ρ* = 0.99), we performed a subject-wise bootstrap, sampling 24 hemispheres from all the 24 hemispheres (12 participant x 2 hem) in the dataset with replacement and estimated the group-level MLE correlation estimate for each new sample (Diedrichsen et al., 2026). Finally, the 95% confidence interval (CI) of the correlation estimates was computed.

### Modeling of bimanual representational structure

To answer which factors, contribute to the representation of bimanual movements, we modelled the cross-validated second-moment matrix of the 36 bimanual conditions. The cross-validated second-moment matrix, i.e. the covariance between patterns without removing the pattern mean, is a complementary representation to the cross-validated Euclidian distance matrix (Eq. 1) that allows for optimal statistical modeling of pattern components (Diedrichsen et al., 2020). We first estimated each element of the second moment matrix (*G_i_*_,j_) using the command `pcm.util.estimate_G_crossval()` in the pcmPy toolbox, which uses the following formula:

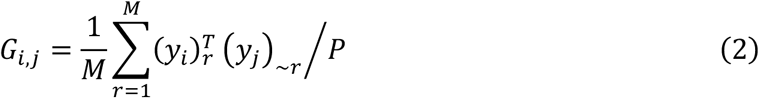

Where (*y*_j_) is the average pattern for condition *j*, across all runs except *r*, and the other variables are the same as in Eq. 1. Because noise is assumed to be independent across runs, this cross-validation (analogous to Eq. 1) ensures that the variance and covariance estimations are not biased due to the shared noise within the run.

For each ROI (or searchlight) in each hemisphere and participant, we then modelled the vectorized second-moment matrix of bimanual conditions (*Ĝ_bimanual_*) as a linear combination of different model components (*w_i_G_i_*):

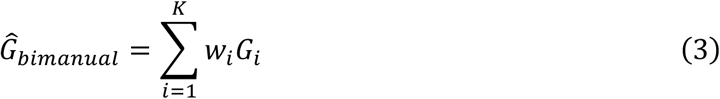

The coefficients (*w_i_*) were fitted by minimizing the squared error between the cross validated second-moment matrix (Eq. 2) and the predicted second moment matrix (Eq. 3). Coefficients were allowed to be both positive and negative to enable us to test them for statistical significance by using a one-sample t-test against 0.

#### Model components

We considered the following model components to explain the structure of the bimanual activity patterns. All components were double centered and normalized by their trace prior to model fitting.

*Contralateral component* assumes that the bimanual structure can be explained by the structure of the contralateral hand activity patterns. To build this component, we estimated the average second-moment matrix from the activity patterns of the six unimanual conditions (using Eq. 2; see supplementary). We then linearly expanded this 6 by 6 matrix to a contralateral 36 by 36 matrix such that the variances and covariances matched those of the unimanual condition in which the contralateral hand was moving to the same target.

*Ipsilateral component* assumes that the bimanual structure can be explained by the structure of the ipsilateral hand activity patterns. This component was built in a way analogous to the contralateral component.

*Bimanual interaction component* assumes that each bimanual combination is uniquely represented. It captures a non-linear modulation of the contralateral by the ipsilateral movement direction. This component was defined as an identity matrix, with all covariances set to 0.

*Congruency component* was built due to the consideration of the bimanual interference effect. It simply assumes that the incongruent bimanual combinations evoke a higher activity than the congruent ones. This was built by defining a 36 by 1 column vector *c*, which codes the congruent combinations as 0 and the incongruent ones as 1. *cc^T^* then produces the 36 by 36 model component.

### Statistical Analyses

In our experiment we symmetrically assessed the directional encoding for both left and right hand in both left and right hemispheres. In the supplementary section we tested for hemispheric asymmetries. Some differences were found for mean activity, but no differences were found for the multi-variate measures of unimanual and bimanual representation. Therefore, for most of the paper we averaged the estimates across the two hemispheres within each hemisphere, yielding statistical tests with 11 dof. Throughout the manuscript, mean values are reported with ± indicating the standard error of the mean across participants.

Tests of whether activation relative to rest, cross-validated distance, model component weight, and intrinsic and extrinsic pattern correlation exceeded zero were one-sided (note that these quantities have an expected value of zero under the null hypothesis). One-sided tests were also used for the following comparisons, where the direction of the effect was predicted *a priori*: greater activation and greater directional encoding for contralateral than for ipsilateral movements; greater activation for bimanual than for unimanual contralateral movements; greater activation for incongruent than for congruent bimanual movements; and a reduction in contralateral and in ipsilateral encoding from the unimanual to the bimanual condition. All other comparisons were two-sided. To account for multiple comparisons across the seven ROIs, p-values were corrected using the Benjamini-Hochberg false discovery rate process and are reported as pfdr. Uncorrected p-values are reported as puncorrected where relevant.

## Results

### Behavioural performance

Participants were instructed to make rapid out-and-back movements with one or both wrists to one of six peripheral targets (Fig. 1A). In the unimanual conditions, the average RT was 519 ± 12 ms and average MT was 666 ± 23 ms. Movement trajectories were relatively straight and directed towards each target (Fig. 1C). To quantify the movement accuracy, we calculated the angular error of the cursor from the target at the peak reaching amplitude. Across participants, the error for the left hand (4.12 ± 0.19 deg) was slightly larger than for the right hand (4.06 ± 0.17 deg); however, this difference was not significant (t_11_ = 0.315, p = 0.759; fig. 1D). In the unimanual condition, participants succeeded in holding the other hand still. The average deviation of the non-moving hand cursor during the trial was 2.1 ± 0.8mm, and the direction of these movements distributed randomly and unrelated to the target of the other hand (fig. 1D). These results indicate that unimanual hand movements did not cause systematic involuntary movements of the other hand.

In the bimanual conditions, participants reached to all possible 36 target pairs (6 left x 6 right; fig. 1C black trajectories). While the RT was not significantly different from the unimanual (523 ± 11 ms, t_11_ = 1.39, p = 0.19), the MT was significantly slower (765 ± 22 ms, t_11_ = 13.97, p = 2.4e-8). We considered three types of bimanual conditions: *intrinsic match*, in which the left and right wrists produce similar movements in joint space (e.g., both flexing inward); *extrinsic match*, in which the left and right targets share the same spatial angle (e.g., both at 60°); and *incongruent*, in which the movements of the two hands were not related intrinsically or extrinsically. The absolute angular error for intrinsic and extrinsic conditions was not significantly different from the unimanual conditions (t_11_ < 0.656, p > 0.525; fig. 1E). However, we observed a slight bimanual interference effect. The absolute angular error during incongruent movements was 0.31 ± 0.05 deg larger than during intrinsically and extrinsically matching movements (both t_11_ > 3.09, p < 0.011). No significant difference was found between intrinsically and extrinsically matching movements (t_11_ = 0.28, p = 0.78).

### Contralateral and ipsilateral motor regions activate during unimanual movements

We first determined the changes in the blood-oxygenation level-dependent (BOLD) signal when participants moved unimanually. As expected, we found signal increases in the primary motor cortex (M1), the somatosensory cortex (S1), in the ventral (PMv) and dorsal (PMd) premotor cortex, in the supplementary motor area (SMA) and in the anterior (SPLa) and posterior (SPLp) superior parietal lobules (fig. 2A). In a Regions of Interest (ROI) analysis (fig. 2B), the increases in the BOLD signal were significant in all these regions during both contralateral and ipsilateral hand movements (all t_11_ > 5.9, p_fdr_ < 5.1e-5).

The activity in PMv was approximately equal during contra- and ipsilateral hand movements (t_11_ = 0.69, p_fdr_ = 0.75). In all other ROIs, the contralateral activity was significantly larger than ipsilateral activity (t_11_ > 2.05, p_fdr_ < 0.037). This difference was especially large in M1 and S1, where the contralateral activity was ∼3.5x larger than ipsilateral activity (fig. 2B).

### Movement direction is encoded in both contralateral and ipsilateral motor regions

Regions that are involved in the control of the wrist movement should not only be activated during the movements but also encode the movement direction of each hand. To measure the strength of this encoding, we estimated the average cross-validated distance between the activity patterns associated with each of the six unimanual movement directions both for the left and right hand separately (fig. 3A; see searchlight analysis in methods).

**Figure 3:**
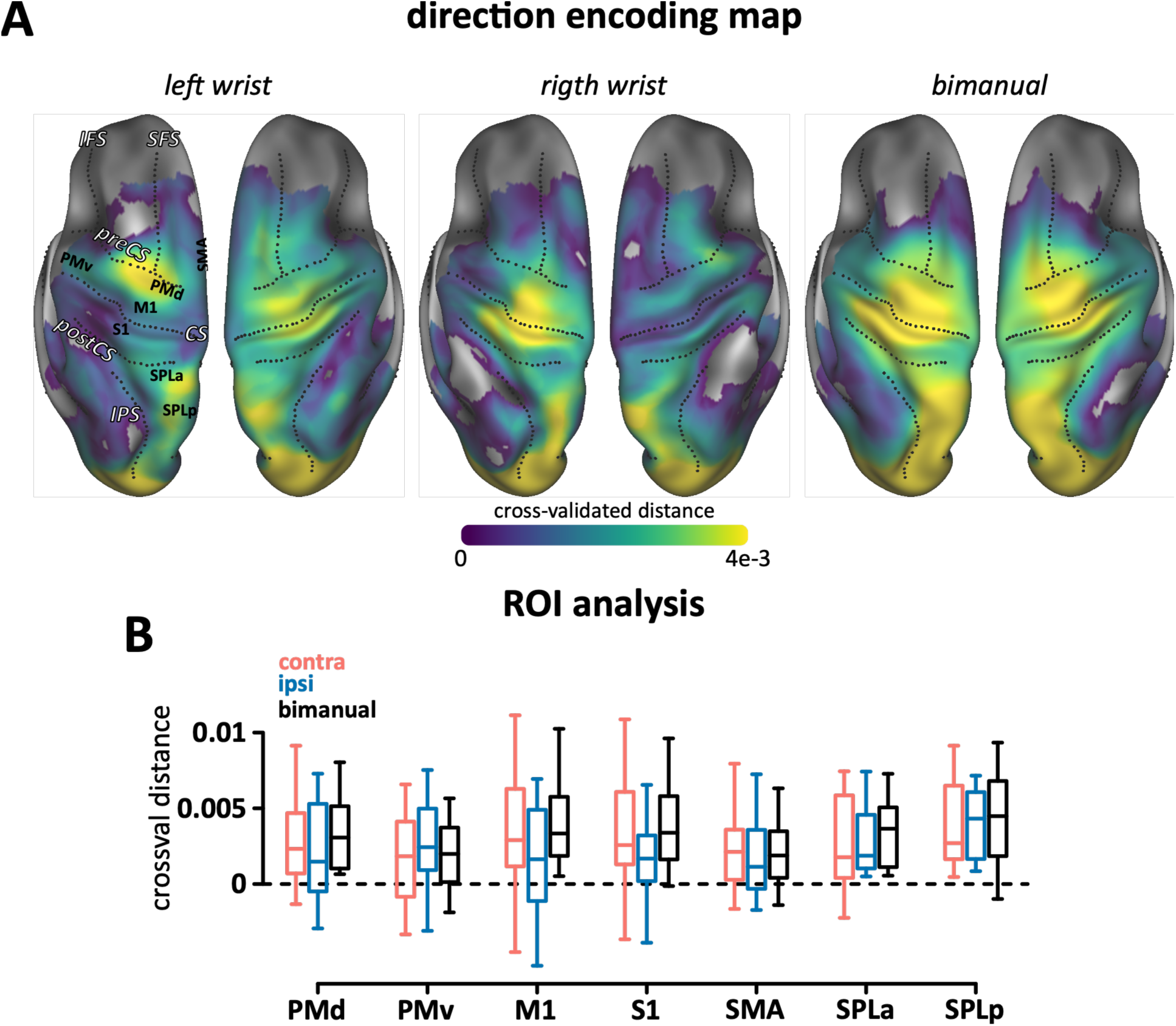
Directional encoding during unimanual and bimanual movements. **A)** Group-averaged cross-validated distance mapped between movement directions obtained from a searchlight analysis across the cortical surface, shown for the left and right hemisphere. Approximate locations of regions of interest (ROIs) are labeled. Direction encoding for the bimanual movements is the average distance between all 36 movement combinations. Maps are thresholded to only show the non-negative cross validated distance values. **B)** Average distance within each ROI for contralateral (red), ipsilateral (blue) and bimanual (black) wrist movements.

Because the dissimilarity measure was cross validated (see methods) we could test the average dissimilarity against zero to determine whether an ROI showed a significant encoding for movement direction (fig. 3B). In all ROIs we found significant direction encoding of the movement of both the contralateral (t_11_ > 2.03, p_fdr_ < 0.033) and the ipsilateral hand (t_11_ > 2.01, p_fdr_ < 0.041) hand, except in M1 in which the ipsilateral encoding was positive but did not reach statistical significance (t_11_ = 1.51, p_uncorrected_ = 0.079). In M1 and S1, the encoding for the contralateral movement was larger than for the ipsilateral movement, although this difference was not significant after correction across the seven ROIs (M1 t_11_=2.01, p_uncorrected_=0.035; S1 t_11_=2.32, p_uncorrected_=0.020; p_fdr_ > 0.12). In all other ROIs, we found no significant differences between the two (t_11_ < 1.62, p_uncorrected_ > 0.067, p_fdr_ > 0.15). These results suggest a relatively bilateral representation of movement direction in parietal and premotor areas, and a strong contralateral representation in primary sensorimotor regions. Nonetheless, encoding of the ipsilateral hand movement was found in all ROIs.

### Ipsi- and contralateral encoding match in intrinsic reference frame

We found that in parietal, motor and premotor regions both the contra- and ipsilateral movement was represented. How does the same population of neurons organize these two distinct movements? We considered three possibilities. First, the activity patterns of contralateral and ipsilateral could be independently organized, such that the patterns indicating movements of the left and right hands are uncorrelated. Neurophysiological studies of M1 have argued for such an orthogonal organization, thereby allowing the regions to represent the movement without causing unintended movement of the non-moving hand (Ames and Churchland, 2019).

Alternatively, the contra- and ipsilateral activity patterns could be either related in the joint/muscle space (body reference frame or intrinsic; fig. 4A: bottom panel) or related in the direction space (world reference frame or extrinsic; fig. 4B: bottom panel).

**Figure 4:**
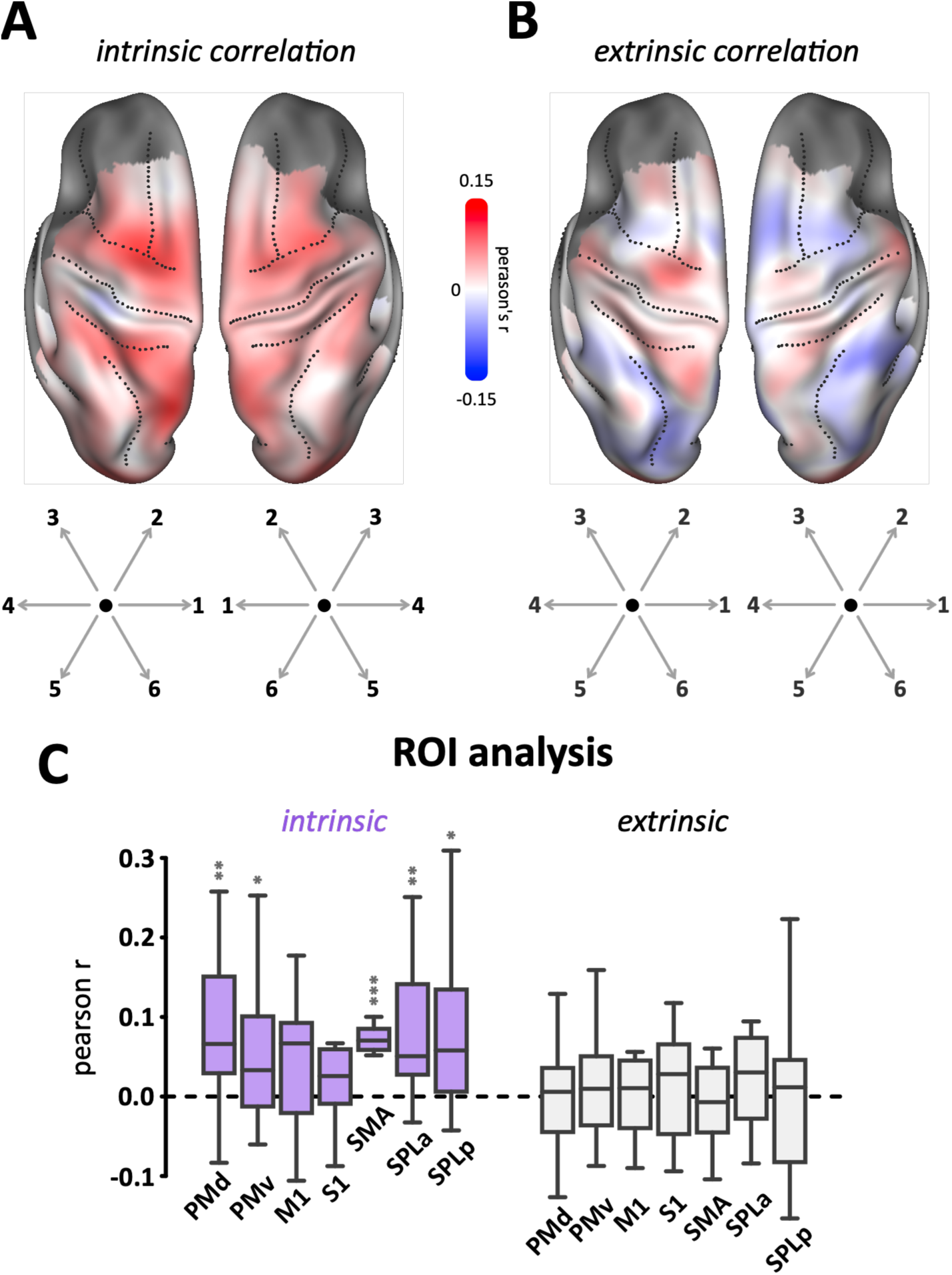
Relationship between contra- and ipsilateral activity is intrinsic. **A)** Pearson’s correlation between brain activity patterns evoked by conditions in which the contralateral and ipsilateral wrists move to similar directions in joint space (intrinsic, e.g., 1: both flex inwards). **B)** Same analysis as in A, but for conditions in which the contralateral and ipsilateral wrists move to the same direction in extrinsic space (e.g., 1: left wrist flex inwards and right wrist extend outwards). **C)** Pearson correlation within each ROI, averaged across hemispheres.

To test this, we correlated the activity patterns for corresponding intrinsic (or extrinsic) movements across the contra- and ipsilateral hand. Intrinsic correlations were positive and significant across premotor and parietal regions (t_11_ > 2.63, p_uncorrecteed_ < 0.012, p_fdr_ < 0.02; fig. 4A,C), although PMv did not survive multiple test correction (t_11_ = 1.93, p_uncorrected_ = 0.04, p_fdr_ = 0.056). In and M1 and S1, however, the correlations were positive but did not reach statistical significance (t_11_ < 1.61, p_uncorrected_ > 0.068, p_fdr_ > 0.08). The extrinsic correlations were not significant in any ROI after multiple tests correction (t_11_ < 1.36, p_uncorrected_ > 0.10, p_fdr_ > 0.48; Fig. 4B,C). Together, these results indicate that most regions exhibit a similar activity pattern for wrist movements that match in an intrinsic (body-centric) reference frame rather than an extrinsic (world-centric) one.

### Bimanual movement activity shows congruency effect

During bimanual movements, we found significant increased BOLD activity relative to rest across all the ROIs (Fig. 2A, B; t_11_ > 8.38, p_uncorrected_ < 2.082e-6 p_fdr_ < 2.082e-6). In M1 and S1, the bimanual activation was only marginally larger (1.06 ± 3.81%) than the contralateral unimanual activation– and this small increase was not significant across participants (both t_11_ < 0.97, p_uncorrected_ > 0.17 p_fdr_>0.20). In the other ROIs however, the activation for bimanual was significantly larger than for contralateral movements (t_11_ > 2.51, p_uncorrected_ < 0.015 p_fdr_ < 0.021).

Based on previous results (Sadato et al., 1997; Debaere et al., 2004; Wenderoth et al., 2004, 2005; Diedrichsen et al., 2006) we expected that premotor and parietal areas would show higher activity during incongruent as compared to intrinsically or extrinsically matching bimanual movements. In M1, S1, SMA and PMv, a repeated measures ANOVA did not show a significant effect of bimanual congruency (F_2,22_ < 2.78, p_uncorrected_ > 0.08 p_fdr_ > 0.14), while in PMd, SPLa and SPLp we found a significant effect (F_2,22_ > 6.46, p_uncorrected_ < 0.0062, p_fdr_ < 0.015). Post-hoc t-tests revealed that PMd, and parietal ROIs were significantly more active during incongruent movements as compared to average of intrinsic (mirror-symmetric) or extrinsic matching movements (t_11_ > 2.77, p_uncorrected_ < 0.009, p_fdr_ < 0.022; Fig. 5). In none of the ROIs did intrinsic and extrinsic combinations differ significantly after multiple test correction (t_11_ < 2.24, p_fdr_ > 0.32). SPLp showed the largest effect (t_11_ = 2.24, p_uncorrected_ = 0.047, p_fdr_ = 0.32) with extrinsic matching combination yielding higher activation than intrinsic combinations.

**Figure 5:**
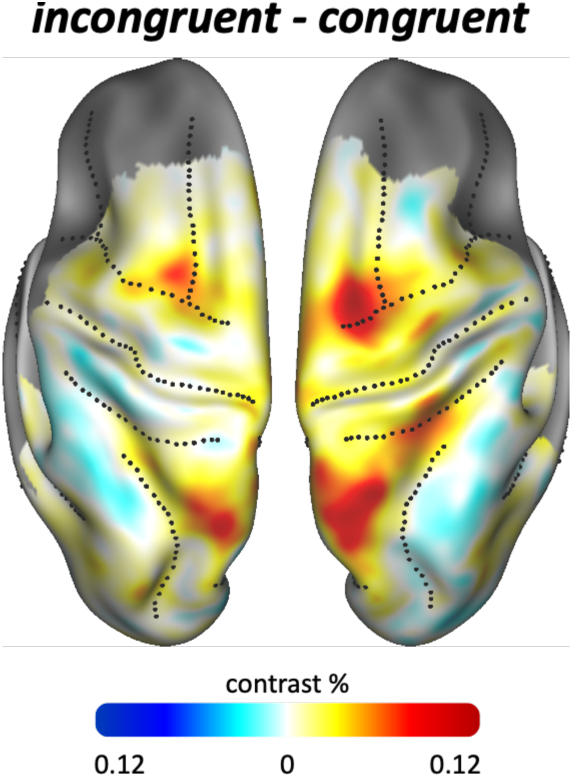
Effect of bimanual incongruency on BOLD activity. Average contrast between the activity of incongruent and congruent bimanual movements.

Overall, the spatial distribution of the congruency effect during bimanual movement (Fig. 5) is largely consistent with previous findings, with the main effect found in posterior parietal areas close to the parietal-occipital sulcus (Diedrichsen et al., 2006). Here we additionally found a clear congruency effect in PMd.

### Representation of contralateral hand movement and bimanual combination

Our main question was how the directional encoding observed during unimanual movements changes when movements are executed bimanually. Using multi-variate analysis, we found that the 36 bimanual conditions were significantly encoded in all the ROIs (Fig. 3A,B; t_11_ > 2.79, p_uncorrected_ < 0.009 p_fdr_ < 0.009). To understand how the different movements were encoded, we built a model that tried to explain the bimanual activity patterns using additive combinations of four components (Fig. 6A): First, we considered two components related to the contralateral movement direction, and the ipsilateral movement direction. These components should be as high as during unimanual movements if the two unimanual patterns simply combined additively. Additionally, we tested for a bimanual interaction effect, which predicted that each of the 36 movement combinations would lead to a unique activity pattern in the area. Finally, we added a congruency component that captured the effects caused by the larger activity for incongruent rather than congruent movement. We fitted this model to the 36×36 second-moment matrix of the bimanual activity patterns to determine how strongly each component is encoded.

**Figure 6:**
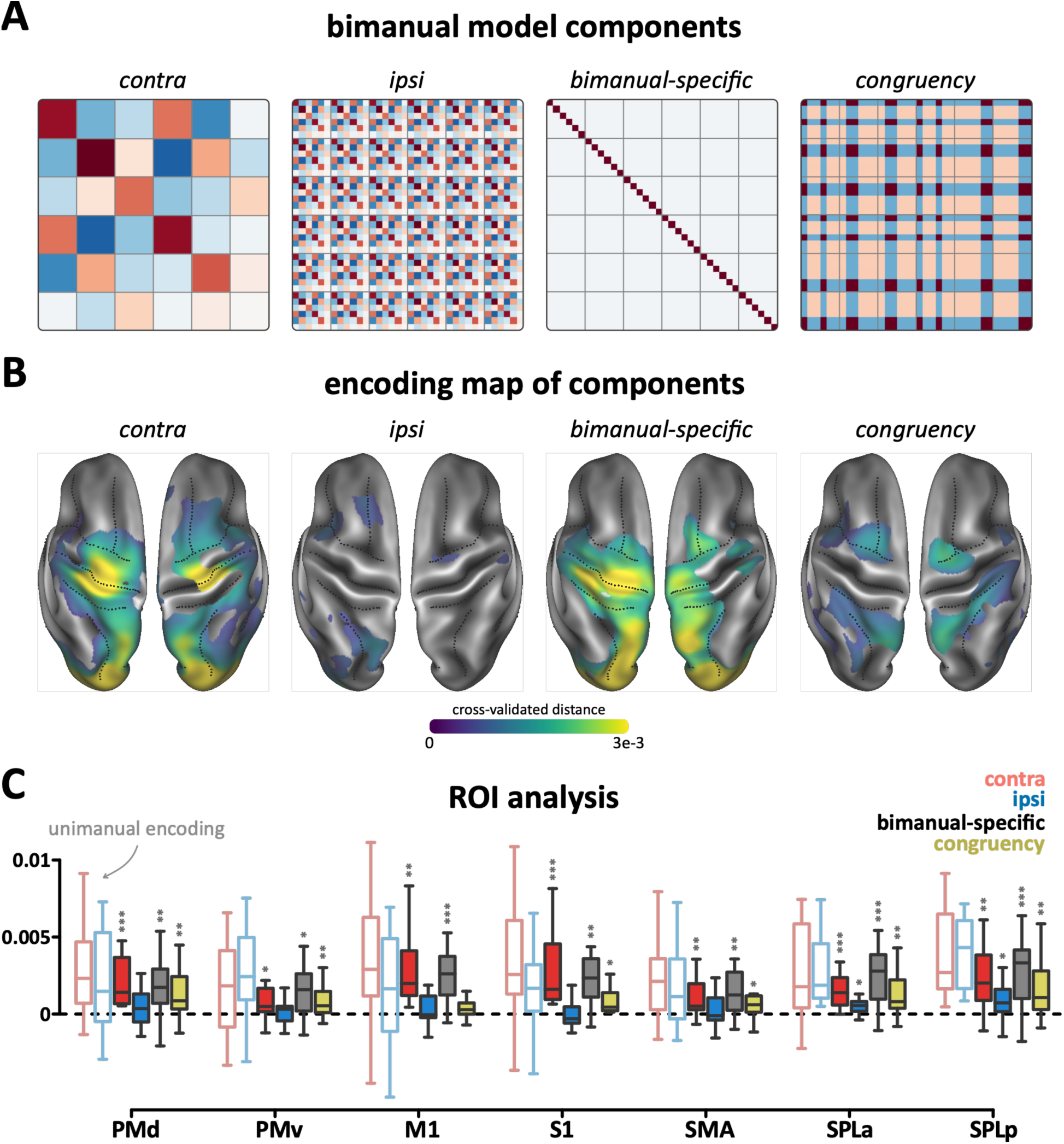
Bimanual activity patterns decomposition. **A)** The brain activity patterns during bimanual conditions decomposed into contralateral, ipsilateral, bimanual interaction, and congruency components. **B)** Searchlight map of component encoding strength expressed as cross-validated Euclidean distance. The model was fitted to the activity patterns of each searchlight to derive their contribution to the overall bimanual movement encoding. Maps are thresholded where distance values are significantly larger than 0; p<0.05. **C)** Component encoding strength within each ROI. The two leftmost boxes in each ROI show the encoding measured in unimanual movements for the contralateral and ipsilateral hand and serve as a reference. The filled boxes show the encoding strength of the components from the model. One-sided t-test > 0; uncorrected p<0.05:*, p<0.01:**, p<0.001:***

The contralateral component encoding was significantly larger than 0 in all ROIs (Fig. 6B, C; t_11_ > 2.10, p_uncorrected_ < 0.03 p_fdr_ < 0.03). To answer whether this component was as strong during bimanual movements as during unimanual movements, we compared the average cross-validated distance between different movement directions for the unimanual conditions (faint boxes Fig. 6B) with the distances between the average activity patterns for the same hand during bimanual movements. Although the bimanual estimate tended to be slightly lower than the unimanual estimate, the difference was not significant in any of the ROIs (t_11_ < 1.76, p_uncorrected_ > 0.054 p_fdr_ > 0.11), except for SPLp (t_11_ = 5.09, p_uncorrected_ = 1.7e-4, p_fdr_ = 0.001). The pattern indicating the contralateral movement direction during bimanual movements also appeared to be identical to the pattern observed when the hand moved in isolation. To show this, we computed patterns related to the movement of the contralateral hand by averaging bimanual activity patterns across all 6 directions of the partner hand, yielding 6 marginal activity patterns from the 36 bimanual conditions. We then estimated the noise-corrected correlation between the marginal unimanual and unimanual activity patterns using group-level maximum likelihood estimation (see methods). Interestingly, in all ROIs, the estimated correlation was at ceiling (mean = 1.0, 95% CI [1.00, 1.00]), indicating near-perfect correlation across all bootstrap samples. This provides evidence that the directional pattern associated with movements of the contralateral hand were also present when both hands moved.

In contrast to the strong and persistent activity patterns related to the movement of the contralateral hand, the patterns related to the movement of the ipsilateral hand nearly disappeared during bimanual movement. The encoding of the ipsilateral component was not significantly larger than 0 in any of the regions after multiple test correction (t_11_ < 2.06, p_fdr_ > 0.148). The parietal regions, SPLa and SPLp, presented the largest ipsilateral effects among the regions (p_uncorrected_ = 0.032 and 0.043, respectively). Compared to the unimanual condition, the ipsilateral also encoding decreased significantly (t_11_ > 2.43, p_uncorrected_ < 0.017 p_fdr_ < 0.019; Fig. 6B, C), except for M1 (t_11_ = 1.53, p_uncorrected_ = 0.077, p_fdr_ = 0.077), where the unimanual ipsilateral encoding failed to become significant. Thus, the ipsilateral activity pattern that existed during unimanual movements mostly disappeared when moving bimanually.

Although we found no evidence for an encoding of the ipsilateral movement independent of the contralateral movement, we found clear evidence that the two movements interacted non-linearly. The bimanual interaction component, which captures any non-additive effect (except for the congruency effect), was significantly larger than 0 in all ROIs (t_11_ > 2.64, p_uncorrected_ < 0.012, p_fdr_ < 0.012; Fig. 6B, C).

The congruency component was significantly larger than zero in six of the seven ROIs: in PMd, PMv, SPLa and SPLp (t_11_ > 2.74, p_uncorrected_ < 0.010, p_fdr_ = 0.017; Fig. 6B, C), in S1 (t_11_ = 2.18, p_uncorrected_ = 0.026, p_fdr_ = 0.037), and in SMA (t_11_ = 1.89, p_uncorrected_ = 0.043, p_fdr_ = 0.050). In M1 however, the congruency component did not reach significance (t_11_ = 1.30, p_uncorrected_ = 0.110, p_fdr_ = 0.110).

For further visualization we also averaged the encoding of each component along the strip running from rostral PMd to caudal SPLp (Fig. 7), providing a profile plot of encoding across the sensorimotor strip. Both visualizations clearly show that the congruency effect was most pronounced in PMd and SPLp, whereas the representational structure of M1, S1 was more dominated by a bimanual interaction code. Correspondingly, the region x weight interaction was significant (F_6,66_ = 3.71, p = 0.003). Post-hoc comparisons revealed significant differences in the primary sensorimotor regions (t_11_ > 3.22, p_uncorrected_ < 0.008, p_fdr_ < 0.029), while the premotor and parietal regions did not reach significance (t_11_ < 2.04, p_uncorrected_ > 0.06, p_fdr_ > 0.12).

**Figure 7:**
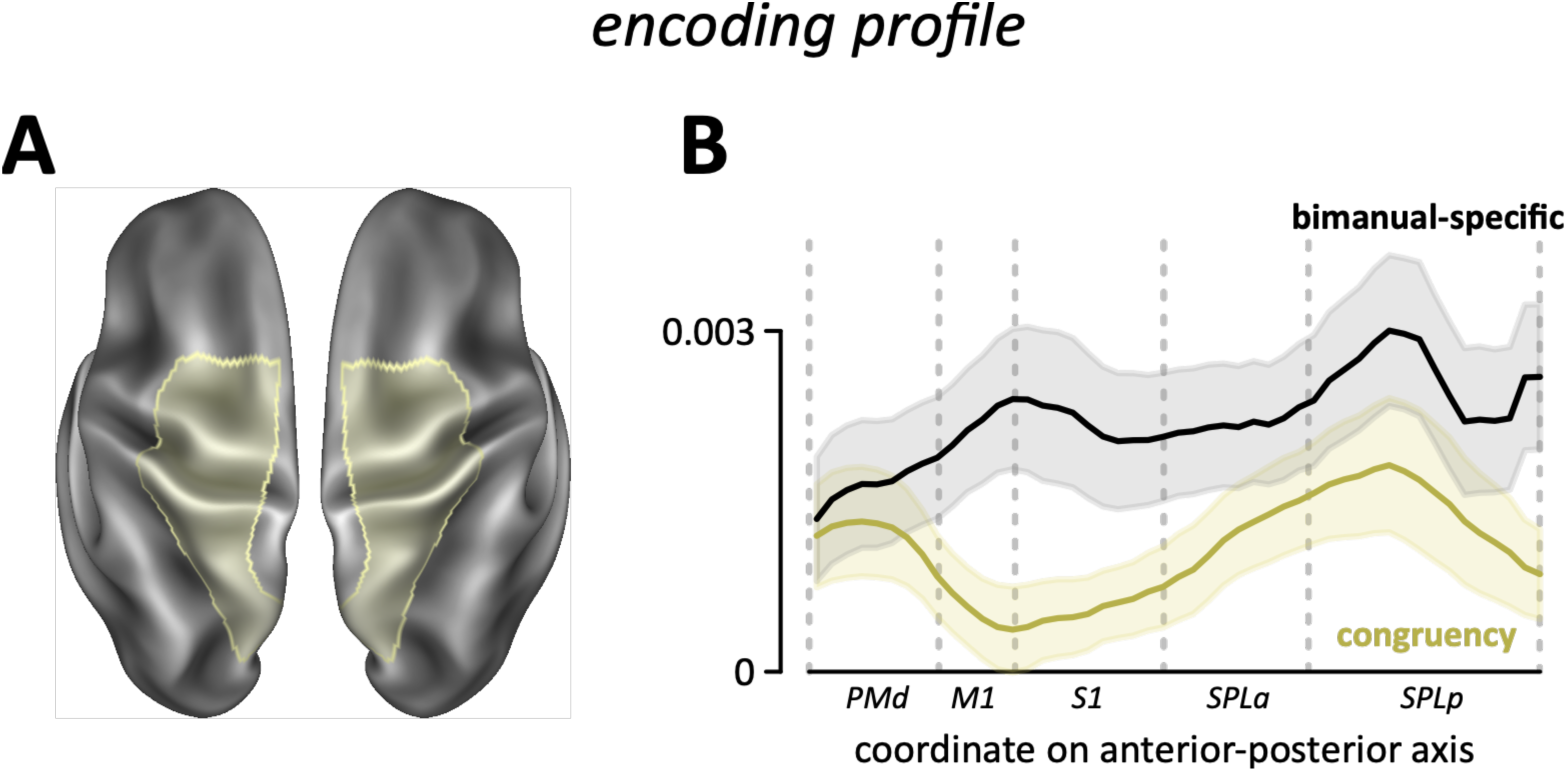
Bimanual-specific encoding vs. congruency effect. **A)** The highlighted area in the anatomical surface map denotes the strip which includes the anterior dorsal premotor to the posterior superior parietal regions. **B)** The encoding profile is the strip average of the encoding from anterior to posterior sensorimotor regions, averaged across the hemispheres. Shaded areas around the lines denote the SEM across participants. Vertical gray lines denote the approximate location of each region.

## Discussion

The primary objective of this study was to determine how bimanual reaching movements are encoded in the human motor system. We hypothesized that the architecture should be able to support relatively independent movements across the two hands, while at the same time providing the computational basis for bimanual interaction (Yokoi et al., 2011). Here we demonstrated that each hemisphere encodes bimanual movements with an activity pattern that relates to the movement of the contralateral hand, which is then non-linearly modulated by the ipsilateral movement, resulting in an interactive code of specific bimanual movement combination. This movement representation enables independent movement of the two hands, and at the same time, the ability to coordinate between the hands.

### Stable and independent encoding of the contralateral hand

Motor, pre-motor, and parietal regions of each hemisphere encoded the movement direction of the contralateral hand in a consistent manner, no matter if the movement was executed in a unimanual or bimanual context. This is consistent with the idea that each hemisphere controls the contralateral hand (Penfield and Boldrey, 1937; Rathelot and Strick, 2006; Soteropoulos et al., 2011), and therefore must produce a stable control signal to the lower spinal and brainstem circuits. Consistent encoding of the contralateral movement independent of whether the ipsilateral hand moved or not, has also been found in electrophysiological recording (Rokni et al., 2003; Mooshagian et al., 2018, 2021; Cross et al., 2020). Here we show that this consistent contralateral component is not restricted to primary sensorimotor areas but also is present in premotor cortex and in superior parietal areas. These stable representations may underly the generalization of force field learning from unimanual to the bimanual movements (Nozaki et al., 2006; Takiyama and Sakai, 2016).

### The evidence for encoding of the specific bimanual combination

The ipsilateral movement modulated the activity patterns of the contralateral movement, resulting in the non-linear encoding of the specific bimanual combination superimposed on stable encoding of the contralateral movement. We use non-linear here referring to any encoding that is not explainable by a linear combination of the patterns related to the unimanual components. Previous behavioural studies had predicted that such a bimanual code must exist in the nervous system (Nozaki et al., 2006; Yokoi et al., 2011, 2014). For example, Yokoi et al. (2011) adapted the left arm of participants to a forcefield, while both left and right arms simultaneously reached forward. The amount of adaptation on the left arm was then probed while the left arm continued moving forward and the right arm moved in different directions. The level of adaptation expressed by the left arm was maximal when the right arm moved forward and fell off for other movement directions. However, even when the right arm moved orthogonal or opposite to its original direction, the left arm still showed some adaptation. This pattern of result is consistent with a model in which the ipsilateral movement direction non-linearly modulated the tuning to the contralateral movement, leading to a bimanual-specific code superimposed on the encoding of the contralateral movement, as observed in our study.

The non-linear bimanual code is also consistent with previous electrophysiological studies in the non-human primate. Single neurons in M1, PMd, SMA, and parietal areas generally respond to both unimanual and bimanual reaching movements (Donchin et al., 1998, 2001, 2002; Steinberg et al., 2002; Rokni et al., 2003; Mooshagian et al., 2018; Cross et al., 2020). The directional tuning of these neurons during the bimanual condition was not a linear combination of the tuning in the respective unimanual conditions. This was the case both in M1 (Rokni et al., 2003) and parietal regions (Mooshagian et al., 2018). As in our current fMRI results, the bimanual tuning in both these areas consisted of a component that reflected the contralateral movement direction, and one that reflected the specific bimanual movement combination.

Our results contrast with findings by Cross et al. (2020), who found that the tuning of M1 neurons during bimanual movements could be explained well by a linear combination of the two unimanual movements. While several factors may explain this discrepancy, the largest difference is that our study, as well as Rokni et al. (2003), used voluntarily initiated movements, whereas Cross et al. (2020) applied mechanical loads to the arm. Thus, their results may reflect that contra- and ipsilateral sensory information combines approximately linearly, whereas the movement plans for simultaneous voluntary movements integrate non-linearly.

Overall, therefore, for voluntary movements, existing studies find a non-linear representation of bimanual movement directions (a uniquely bimanual code) in primary sesnorimotor (M1 and S1), premotor and parietal regions. This is also consistent with our previous findings for bimanual finger movements (Diedrichsen et al., 2013).

### The bimanual congruency effect

In addition to a non-linear encoding of the bimanual movement combination, we found a distinct third component, reflecting the higher activation during incongruent bimanual movements. This movement congruency effect was mostly absent from the primary sensorimotor regions and instead most pronounced in rostral premotor and caudal parietal regions. In contrast, the bimanual movement encoding was more evenly distributed across the entire hierarchy of motor and premotor regions. This spatial distinction suggests that the bimanual and congruency components are fundamentally different.

The rostral premotor and caudal parietal regions are involved in abstract sensory representation (Gallivan et al., 2011), movement preparation (Ariani et al., 2022, 2025), and visual attention (Jerde et al., 2012). Previous imaging work consistently reports higher activation in these regions during bimanual interference tasks (Debaere et al., 2004; Wenderoth et al., 2004, 2005; Diedrichsen et al., 2006). This localization aligns with behavioural evidence for the perceptual basis of bimanual interference (Mechsner et al., 2001). For example, bimanual interference is not only reduced when participants move in a mirror-symmetric fashion engaging homologous muscle groups, but also when the cue and resultant movement go in the same direction in extrinsic space (Mechsner et al., 2001). Therefore, our results are consistent with the idea that bimanual interference arises at more abstract levels of the motor hierarchy (goal selection, stimulus-response mapping) rather than from the interaction of motor-related processes in M1 and S1 (Diedrichsen et al., 2001, 2003a, 2006).

### Ipsilateral movement representation

Our findings suggest that each hemisphere encodes bimanual movements as a combination of the contralateral movement and the specific bimanual movement combination, without any linearly separable code for the ipsilateral movement. In contrast, we found a clear representation of ipsilateral movement in the unimanual condition, a representation that has been also shown in many electrophysiological studies (Tanji et al., 1988; Donchin et al., 1998; Cisek et al., 2003; Ganguly et al., 2009; Ames and Churchland, 2019; Heming et al., 2019; Cross et al., 2020).

Our results also show that the contralateral and ipsilateral movement representations within the premotor and parietal regions of a hemisphere are not independent from each other but rather are similar for mirror-symmetric movements. For example, flexing the left wrist to the right, evokes an activity pattern similar to flexing the right wrist toward to the left. In contrast, movements in the same spatial direction are not coded in a similar way. This aligns with earlier fMRI and intra-cranial recordings (Diedrichsen et al., 2013; Haar et al., 2017; Zhang et al., 2017; Willett et al., 2020) showing similar activity patterns for mirror-symmetric movements.

We found that this correlation was not reliable in the primary sensorimotor regions, which is consistent with the electrophysiological studies that find largely independent contralateral and ipsilateral representations during arm cycling (Ames and Churchland, 2019) and load application to the arm (Heming et al., 2019). In another study, Downey et al. (2020) recorded from the human motor cortex using intracortical microelectrode arrays and reported high correlation between contralateral and ipsilateral grasping movements with little to no correlation between contralateral and ipsilateral arm directional reaching. This suggests that this contra-ipsi relationship might depend on the effector and the task that is performed.

The observation that ipsilateral movement encoding is not fully independent from the contralateral movement during unimanual movements, and then disappears as an linearly-separable component during bimanual movements, raises the question of its function (Berlot et al., 2019). One possibility is that each hemisphere is causally involved in the control of the ipsilateral hand, either directly through the small fraction of uncrossed cortico-spinal projections (Dum and Strick, 1996; Rosenzweig et al., 2009) or indirectly by modulating activity in the other hemisphere through callosal projections. While this remains a possibility, a direct role in control must be ultimately relatively restricted, as patients with unilateral lesions to primary motor cortex or the cortical spinal tract show only mild deficits in the ipsilateral limb (Jones et al., 1989; Sunderland, 2000; Semrau et al., 2017; Maenza et al., 2020). These deficits are mainly associated with the coordination, sequencing and dexterity rather than the capacity to move (Yarosh et al., 2004; Schaefer et al., 2009).

Alternatively, therefore, the tight communication between the two motor cortices may mostly have evolved to support bimanual coordination, such that the ipsilateral movement can modulate the contralateral movement in a goal-directed manner. During unimanual movements, the same connections cause neural activity in the other hemisphere. The resulting ipsilateral representation could therefore be either epiphenomenal without a functional role, or it could subserve intermanual transfer of learning (Berlot et al., 2019).

### Hemispheric asymmetries

In this paper, we analyzed the data from the left and right hemisphere together. Qualitatively, we found similar results in both hemispheres. However, there were subtle differences between the left (dominant) and right (non-dominant) hemispheres (see supplementary material). In summary, contralateral movements elicited comparable activity across hemispheres. Ipsilateral movements elicited more activity in the left hemisphere and bimanual movements elicited more activity in the right hemisphere. Notably, these asymmetries varied across the ROIs. These findings partially align with existing literature, which generally reports a left-hemisphere predominance in right-handers across contralateral, ipsilateral (Haaland and Harrington, 1996; Dassonville et al., 1997; Ziemann and Hallett, 2001; Hayashi et al., 2008; Callaert et al., 2011; Tzourio-Mazoyer et al., 2015) and bimanual movements (Viviani et al., 1998; Serrien et al., 2006), despite some mixed results (Hayashi et al., 2008; Schweisfurth et al., 2018).

Importantly, however, consistent with recent work (Schweisfurth et al., 2018; Merrick et al., 2022) we found no significant hemispheric asymmetries in encoding strength, nor in the way that contralateral and ipsilateral movements were encoded. Overall, these findings suggest that while asymmetries in overall activation exist, the actual information encoding is balanced. Thus, functional interpretations drawn solely from mean activation asymmetries should be approached with caution.

## Conclusion

Our study provides a careful multivariate analysis of fMRI activity during unimanual and bimanual reaching movements in humans. Similar to earlier results for the control of finger movements (Diedrichsen et al., 2013), we found encoding of both the contralateral and the ipsilateral movement directions across most sensorimotor, premotor, and superior parietal regions, with the two representations aligned in a mirror-symmetric reference frame in premotor and parietal regions. When these two movements were performed simultaneously in the bimanual conditions, the encoding of the contralateral movement remained unchanged, i.e., the directional pattern was statistically indistinguishable from the one measured in unimanual condition. The ipsilateral movement, however, was no longer encoded as an independent component. Instead, the two movements interacted non-linearly, such that each region carried a unique representation of the specific bimanual combination. This bimanual-specific code was above over and above the increase in activity caused by moving the hands incongruently. Such a code provides what between-hand coordination requires: a stable command for each hand that can be adjusted based on what the other hand is doing.

## Data and Code Availability

Analysis code used in this study is available at https://github.com/alighavam/bimanual_wrist. Derived data supporting the findings – behaviour, ROI activity patterns, and group-level surface maps – will be available in a public repository upon publication. Raw imaging data are available from the corresponding author upon reasonable request, subject to the data-sharing provisions of the ethics approvals at each collection site.

## Acknowledgements

Present affiliations: A. Yokoi, Center for Information and Neural Networks (CiNet), National Institute of Information and Communications Technology (NICT), Osaka, Japan. D. F. Duarte, Digital Data Design Institute, Nova School of Business and Economics, Carcavelos, Portugal.

## Author Contributions

<u>Ali Ghavampour</u>: Conceptualization, Data Acquisition, Methodology, Analysis, Writing – original draft, review and editing; <u>Atsushi Yokoi</u>: Conceptualization, Data Acquisition, Methodology, Analysis, Writing – review and editing; <u>Diogo F. Duarte</u>: Conceptualization, Methodology, Writing – review and editing; <u>Jean-Jacques Orban de Xivry</u>: Conceptualization, Supervision, Methodology, Writing – review and editing; J<u>. Andrew Pruszynski</u>: Conceptualization, Supervision, Methodology, Writing – review and editing; <u>Jörn Diedrichsen</u>: Conceptualization, Supervision, Methodology, Writing – review and editing;

## References

Albert NB, Ivry RB (2009) The persistence of spatial interference after extended training in a bimanual drawing task. Cortex 45:377–385. doi: 10.1016/j.cortex.2007.11.012.

Ames KC, Churchland MM (2019) Motor cortex signals for each arm are mixed across hemispheres and neurons yet partitioned within the population response. Elife 8. doi: 10.7554/eLife.46159.

Arbuckle SA, Yokoi A, Pruszynski JA, Diedrichsen J (2019) Stability of representational geometry across a wide range of fMRI activity levels. Neuroimage 186:155–163. doi: 10.1016/j.neuroimage.2018.11.002.

Ariani G, Pruszynski JA, Diedrichsen J (2022) Motor planning brings human primary somatosensory cortex into action-specific preparatory states. Elife 11. doi: 10.7554/eLife.69517.

Ariani G, Shahbazi M, Diedrichsen J (2025) Cortical Areas for Planning Sequences before and during Movement. J Neurosci 45:e1300242024. doi: 10.1523/JNEUROSCI.1300-24.2024.

Berlot E, Popp NJ, Diedrichsen J (2020) A critical re-evaluation of fMRI signatures of motor sequence learning. Elife 9. doi: 10.7554/eLife.55241.

Berlot E, Prichard G, O’Reilly J, Ejaz N, Diedrichsen J (2019) Ipsilateral finger representations in the sensorimotor cortex are driven by active movement processes, not passive sensory input. J Neurophysiol 121:418–426. doi: 10.1152/jn.00439.2018.

Blinch J, Cameron BD, Cressman EK, Franks IM, Carpenter MG, Chua R (2014) Comparing movement preparation of unimanual, bimanual symmetric, and bimanual asymmetric movements. Exp Brain Res 232:947–955. doi: 10.1007/s00221-013-3807-7.

Callaert DV, Vercauteren K, Peeters R, Tam F, Graham S, Swinnen SP, Sunaert S, Wenderoth N (2011) Hemispheric asymmetries of motor versus nonmotor processes during (visuo)motor control. Hum Brain Mapp 32:1311–1329. doi: 10.1002/hbm.21110.

Cisek P, Crammond DJ, Kalaska JF (2003) Neural activity in primary motor and dorsal premotor cortex in reaching tasks with the contralateral versus ipsilateral arm. J Neurophysiol 89:922–942. doi: 10.1152/jn.00607.2002.

Crisco JJ, Heard WMR, Rich RR, Paller DJ, Wolfe SW (2011) The mechanical axes of the wrist are oriented obliquely to the anatomical axes. J Bone Joint Surg Am 93:169–177. doi: 10.2106/JBJS.I.01222.

Cross KP, Heming EA, Cook DJ, Scott SH (2020) Maintained representations of the ipsilateral and contralateral limbs during bimanual control in primary motor cortex. J Neurosci 40:6732–6747. doi: 10.1523/JNEUROSCI.0730-20.2020.

Dale AM, Fischl B, Sereno MI (1999) Cortical surface-based analysis. I. Segmentation and surface reconstruction. Neuroimage 9:179–194. doi: 10.1006/nimg.1998.0395.

Dassonville P, Zhu XH, Uurbil K, Kim SG, Ashe J (1997) Functional activation in motor cortex reflects the direction and the degree of handedness. Proc Natl Acad Sci U S A 94:14015–14018. doi: 10.1073/pnas.94.25.14015.

Debaere F, Wenderoth N, Sunaert S, Van Hecke P, Swinnen SP (2004) Cerebellar and premotor function in bimanual coordination: parametric neural responses to spatiotemporal complexity and cycling frequency. Neuroimage 21:1416–1427. doi: 10.1016/j.neuroimage.2003.12.011.

Desrochers PC, Brunfeldt AT, Kagerer FA (2026) Visuomotor information drives interference between the hands more than dynamic motor information during bimanual reaching. Exp Brain Res 244:72. doi: 10.1007/s00221-026-07270-5.

Diedrichsen J, Berlot E, Mur M, Schütt HH, Shahbazi M, Kriegeskorte N (2020) Comparing representational geometries using whitened unbiased-distance-matrix similarity. arXiv [statAP]. doi: 10.48550/arXiv.2007.02789.

Diedrichsen J, Fu X, Shahbazi M, Bonner S (2026) Testing hypotheses about correlations between brain activation patterns. *bioRxiv*:2026.03.21.713393.doi: 10.64898/2026.03.21.713393.

Diedrichsen J, Grafton S, Albert N, Hazeltine E, Ivry RB (2006) Goal-selection and movement-related conflict during bimanual reaching movements. Cereb Cortex 16:1729–1738. doi: 10.1093/cercor/bhj108.

Diedrichsen J, Hazeltine E, Kennerley S, Ivry RBRB (2001) Moving to directly cued locations abolishes spatial interference during bimanual actions. Psychol Sci 12:493–498. doi: 10.1111/1467-9280.00391.

Diedrichsen J, Ivry RBB, Kennerley S, Hazeltine E, Cohen A, Kennerley S, Cohen A (2003a) Bimanual interference associated with the selection of target locations. J Exp Psychol Hum Percept Perform 29:64–77. doi: 10.1037/0096-1523.29.1.64.

Diedrichsen J, Verstynen T, Hon A, Lehman SL, Ivry RB (2003b) Anticipatory adjustments in the unloading task: is an efference copy necessary for learning? Exp Brain Res 148:272–276. doi: 10.1007/s00221-002-1318-z.

Diedrichsen J, Verstynen T, Lehman SL, Ivry RB (2005) Cerebellar involvement in anticipating the consequences of self-produced actions during bimanual movements. J Neurophysiol 93:801–812. doi: 10.1152/jn.00662.2004.

Diedrichsen J, Wiestler T, Krakauer JW (2013) Two distinct ipsilateral cortical representations for individuated finger movements. Cereb Cortex 23:1362–1377. doi: 10.1093/cercor/bhs120.

Diedrichsen J, Yokoi A, Arbuckle SA (2018) Pattern component modeling: A flexible approach for understanding the representational structure of brain activity patterns. Neuroimage 180:119–133. doi: 10.1016/j.neuroimage.2017.08.051.

Donchin O, Gribova A, Steinberg O, Bergman H, Cardoso de Oliveira S, Vaadia E (2001) Local field potentials related to bimanual movements in the primary and supplementary motor cortices. Exp Brain Res 140:46–55. doi: 10.1007/s002210100784.

Donchin O, Gribova A, Steinberg O, Bergman H, Vaadia E (1998) Primary motor cortex is involved in bimanual coordination. Nature 395:274–278. doi: 10.1038/26220.

Donchin O, Gribova A, Steinberg O, Mitz AR, Bergman H, Vaadia E (2002) Single-unit activity related to bimanual arm movements in the primary and supplementary motor cortices. J Neurophysiol 88:3498–3517. doi: 10.1152/jn.00335.2001.

Downey JE, Quick KM, Schwed N, Weiss JM, Wittenberg GF, Boninger ML, Collinger JL (2020) The motor cortex has independent representations for ipsilateral and contralateral arm movements but correlated representations for grasping. Cereb Cortex 30:5400–5409. doi: 10.1093/cercor/bhaa120.

Dum RP, Strick PL (1996) Spinal cord terminations of the medial wall motor areas in macaque monkeys. J Neurosci 16:6513–6525. doi: 10.1523/JNEUROSCI.16-20-06513.1996.

Fischl B, Rajendran N, Busa E, Augustinack J, Hinds O, Yeo BTT, Mohlberg H, Amunts K, Zilles K (2008) Cortical folding patterns and predicting cytoarchitecture. Cereb Cortex 18:1973–1980. doi: 10.1093/cercor/bhm225.

Fischl B, Sereno MI, Dale AM (1999a) Cortical surface-based analysis. II: Inflation, flattening, and a surface-based coordinate system. Neuroimage 9:195–207. doi: 10.1006/nimg.1998.0396.

Fischl B, Sereno MI, Tootell RB, Dale AM (1999b) High-resolution intersubject averaging and a coordinate system for the cortical surface. Hum Brain Mapp 8:272–284. doi: 10.1002/(sici)1097-0193(1999)8:4<272::aid-hbm10>3.0.co;2-4.

Franz EA, Zelaznik HN, McCabe G (1991) Spatial topological constraints in a bimanual task. Acta Psychol (Amst) 77:137–151. doi: 10.1016/0001-6918(91)90028-X.

Friston KJ, Jezzard P, Turner R (1994) Analysis of functional MRI time-series. Hum Brain Mapp 1:153–171. doi: 10.1002/hbm.460010207.

Gallivan JP, McLean DA, Smith FW, Culham JC (2011) Decoding effector-dependent and effector-independent movement intentions from human parieto-frontal brain activity. J Neurosci 31:17149–17168. doi: 10.1523/JNEUROSCI.1058-11.2011.

Ganguly K, Secundo L, Ranade G, Orsborn A, Chang EF, Dimitrov DF, Wallis JD, Barbaro NM, Knight RT, Carmena JM (2009) Cortical representation of ipsilateral arm movements in monkey and man. J Neurosci 29:12948–12956. doi: 10.1523/JNEUROSCI.2471-09.2009.

Guan C, Aflalo T, Kadlec K, Gámez de Leon J, Rosario E, Bari A, Pouratian N, Andersen R (2022) Compositional coding of individual finger movements in human posterior parietal cortex and motor cortex enables ten-finger decoding. medRxiv:2022.12.07.22283227. doi: 10.1101/2022.12.07.22283227.

Haaland KY, Harrington DL (1996) Hemispheric asymmetry of movement. Curr Opin Neurobiol 6:796–800. doi: 10.1016/s0959-4388(96)80030-4.

Haar S, Dinstein I, Shelef I, Donchin O (2017) Effector-invariant movement encoding in the human motor system. J Neurosci 37:9054–9063. doi: 10.1523/JNEUROSCI.1663-17.2017.

Hayashi MJ, Saito DN, Aramaki Y, Asai T, Fujibayashi Y, Sadato N (2008) Hemispheric asymmetry of frequency-dependent suppression in the ipsilateral primary motor cortex during finger movement: a functional magnetic resonance imaging study. Cereb Cortex 18:2932–2940. doi: 10.1093/cercor/bhn053.

Heming EA, Cross KP, Takei T, Cook DJ, Scott SH (2019) Independent representations of ipsilateral and contralateral limbs in primary motor cortex. Elife 8. doi: 10.7554/eLife.48190.

Heuer H, Spijkers W, Kleinsorge T, van der Loo H, Steglich C (1998) The time course of cross-talk during the simultaneous specification of bimanual movement amplitudes. Exp Brain Res 118:381–392. doi: 10.1007/s002210050292.

Jerde TA, Merriam EP, Riggall AC, Hedges JH, Curtis CE (2012) Prioritized maps of space in human frontoparietal cortex. J Neurosci 32:17382–17390. doi: 10.1523/JNEUROSCI.3810-12.2012.

Jones RD, Donaldson IM, Parkin PJ (1989) Impairment and recovery of ipsilateral sensory-motor function following unilateral cerebral infarction. Brain 112 (Pt 1):113–132. doi: 10.1093/brain/112.1.113.

Kelso JA (1984) Phase transitions and critical behavior in human bimanual coordination. Am J Physiol 246:R1000–4. doi: 10.1152/ajpregu.1984.246.6.R1000.

Kelso JA, Southard DL, Goodman D (1979) On the nature of human interlimb coordination. Science 203:1029–1031. doi: 10.1126/science.424729.

Kornysheva K, Sierk A, Diedrichsen J (2013) Interaction of temporal and ordinal representations in movement sequences. J Neurophysiol 109:1416–1424. doi: 10.1152/jn.00509.2012.

Maenza C, Good DC, Winstein CJ, Wagstaff DA, Sainburg RL (2020) Functional deficits in the less-impaired arm of stroke survivors depend on hemisphere of damage and extent of paretic arm impairment. Neurorehabil Neural Repair 34:39–50. doi: 10.1177/1545968319875951.

Marcus DS, Harwell J, Olsen T, Hodge M, Glasser MF, Prior F, Jenkinson M, Laumann T, Curtiss SW, Van Essen DC (2011) Informatics and data mining tools and strategies for the human connectome project. Front Neuroinform 5:4. doi: 10.3389/fninf.2011.00004.

Marteniuk RG, MacKenzie CL, Baba DM (1984) Bimanual Movement Control: Information processing and Interaction Effects. Q J Exp Psychol A 36:335–365. doi: 10.1080/14640748408402163.

Mechsner F, Kerzel D, Knoblich G, Prinz W (2001) Perceptual basis of bimanual coordination. Nature 414:69–73. doi: 10.1038/35102060.

Merrick CM, Dixon TC, Breska A, Lin J, Chang EF, King-Stephens D, Laxer KD, Weber PB, Carmena J, Thomas Knight R, Ivry RB (2022) Left hemisphere dominance for bilateral kinematic encoding in the human brain. Elife 11. doi: 10.7554/eLife.69977.

Mooshagian E, Holmes CD, Snyder LH (2021) Local field potentials in the parietal reach region reveal mechanisms of bimanual coordination. Nat Commun 12:2514. doi: 10.1038/s41467-021-22701-3.

Mooshagian E, Wang C, Holmes CD, Snyder LH (2018) Single units in the posterior parietal cortex encode patterns of bimanual coordination. Cereb Cortex 28:1549–1567. doi: 10.1093/cercor/bhx052.

Nozaki D, Kurtzer I, Scott SH (2006) Limited transfer of learning between unimanual and bimanual skills within the same limb. Nat Neurosci 9:1364–1366. doi: 10.1038/nn1785.

Omrani M, Diedrichsen J, Scott SH (2013) Rapid feedback corrections during a bimanual postural task. J Neurophysiol 109:147–161. doi: 10.1152/jn.00669.2011.

Oosterhof NN, Wiestler T, Downing PE, Diedrichsen J (2011) A comparison of volume-based and surface-based multi-voxel pattern analysis. Neuroimage 56:593–600. doi: 10.1016/j.neuroimage.2010.04.270.

Orschiedt J, Franklin DW (2023) Learning context shapes bimanual control strategy and generalization of novel dynamics. PLoS Comput Biol 19:e1011189. doi: 10.1371/journal.pcbi.1011189.

Penfield W, Boldrey E (1937) Somatic motor and sensory representation in the cerebral cortex of man as studied by electrical stimulation. Brain 60:389–443. doi: 10.1093/brain/60.4.389.

Rathelot J-A, Strick PL (2006) Muscle representation in the macaque motor cortex: an anatomical perspective. Proc Natl Acad Sci U S A 103:8257–8262. doi: 10.1073/pnas.0602933103.

Rokni U, Steinberg O, Vaadia E, Sompolinsky H (2003) Cortical representation of bimanual movements. J Neurosci 23:11577–11586. doi: 10.1523/JNEUROSCI.23-37-11577.2003.

Rosenzweig ES, Brock JH, Culbertson MD, Lu P, Moseanko R, Edgerton VR, Havton LA, Tuszynski MH (2009) Extensive spinal decussation and bilateral termination of cervical corticospinal projections in rhesus monkeys. J Comp Neurol 513:151–163. doi: 10.1002/cne.21940.

Sadato N, Yonekura Y, Waki A, Yamada H, Ishii Y (1997) Role of the supplementary motor area and the right premotor cortex in the coordination of bimanual finger movements. J Neurosci 17:9667–9674. doi: 10.1523/JNEUROSCI.17-24-09667.1997.

Schaefer SY, Haaland KY, Sainburg RL (2009) Hemispheric specialization and functional impact of ipsilesional deficits in movement coordination and accuracy. Neuropsychologia 47:2953– 2966. doi: 10.1016/j.neuropsychologia.2009.06.025.

Schweisfurth MA, Frahm J, Farina D, Schweizer R (2018) Comparison of fMRI digit representations of the dominant and non-dominant hand in the human primary somatosensory cortex. Front Hum Neurosci 12:492. doi: 10.3389/fnhum.2018.00492.

Semrau JA, Herter TM, Kenzie JM, Findlater SE, Scott SH, Dukelow SP (2017) Robotic characterization of ipsilesional motor function in subacute stroke. Neurorehabil Neural Repair 31:571–582. doi: 10.1177/1545968317704903.

Serrien DJ, Ivry RB, Swinnen SP (2006) Dynamics of hemispheric specialization and integration in the context of motor control. Nat Rev Neurosci 7:160–166. doi: 10.1038/nrn1849.

Soteropoulos DS, Edgley SA, Baker SN (2011) Lack of evidence for direct corticospinal contributions to control of the ipsilateral forelimb in monkey. J Neurosci 31:11208–11219. doi: 10.1523/JNEUROSCI.0257-11.2011.

Spearman C (1904) The proof and measurement of association between two things. Am J Psychol 15:72. doi: 10.2307/1412159.

Steinberg O, Donchin O, Gribova A, Cardosa de Oliveira S, Bergman H, Vaadia E (2002) Neuronal populations in primary motor cortex encode bimanual arm movements: Population vectors in bimanual movements. Eur J Neurosci 15:1371–1380. doi: 10.1046/j.1460-9568.2002.01968.x.

Sunderland A (2000) Recovery of ipsilateral dexterity after stroke. Stroke 31:430–433. doi: 10.1161/01.STR.31.2.430.

Swinnen SP, Dounskaia N, Duysens J (2002) Patterns of bimanual interference reveal movement encoding within a radial egocentric reference frame. J Cogn Neurosci 14:463–471. doi: 10.1162/089892902317361976.

Swinnen SP, Dounskaia N, Levin O, Duysens J (2001) Constraints during bimanual coordination: the role of direction in relation to amplitude and force requirements. Behav Brain Res 123:201–218. doi: 10.1016/S0166-4328(01)00210-8.

Swinnen SP, Jardin K, Verschueren S, Meulenbroek R, Franz L, Dounskaia N, Walter CB (1998) Exploring interlimb constraints during bimanual graphic performance: effects of muscle grouping and direction. Behav Brain Res 90:79–87. doi: 10.1016/S0166-4328(97)00083-1.

Swinnen SP, Wenderoth N (2004) Two hands, one brain: cognitive neuroscience of bimanual skill. Trends Cogn Sci 8:18–25. doi: 10.1016/j.tics.2003.10.017.

Takiyama K, Sakai Y (2016) Balanced motor primitive can explain generalization of motor learning effects between unimanual and bimanual movements. Sci Rep 6:23331. doi: 10.1038/srep23331.

Tanji J, Okano K, Sato KC (1988) Neuronal activity in cortical motor areas related to ipsilateral, contralateral, and bilateral digit movements of the monkey. J Neurophysiol 60:325–343. doi: 10.1152/jn.1988.60.1.325.

Tzourio-Mazoyer N, Petit L, Zago L, Crivello F, Vinuesa N, Joliot M, Jobard G, Mellet E, Mazoyer B (2015) Between-hand difference in ipsilateral deactivation is associated with hand lateralization: fMRI mapping of 284 volunteers balanced for handedness. Front Hum Neurosci 9:5. doi: 10.3389/fnhum.2015.00005.

Van Essen DC, Glasser MF, Dierker DL, Harwell J, Coalson T (2012) Parcellations and hemispheric asymmetries of human cerebral cortex analyzed on surface-based atlases. Cereb Cortex 22:2241–2262. doi: 10.1093/cercor/bhr291.

Viviani P, Perani D, Grassi F, Bettinardi V, Fazio F (1998) Hemispheric asymmetries and bimanual asynchrony in left- and right-handers. Exp Brain Res 120:531–536. doi: 10.1007/s002210050428.

Walther A, Nili H, Ejaz N, Alink A, Kriegeskorte N, Diedrichsen J (2016) Reliability of dissimilarity measures for multi-voxel pattern analysis. Neuroimage 137:188–200. doi: 10.1016/j.neuroimage.2015.12.012.

Wenderoth N, Debaere F, Sunaert S, Swinnen SP (2005) Spatial interference during bimanual coordination: differential brain networks associated with control of movement amplitude and direction. Hum Brain Mapp 26:286–300. doi: 10.1002/hbm.20151.

Wenderoth N, Debaere F, Sunaert S, van Hecke P, Swinnen SP (2004) Parieto-premotor areas mediate directional interference during bimanual movements. Cereb Cortex 14:1153–1163. doi: 10.1093/cercor/bhh075.

Wiestler T, Diedrichsen J (2013) Skill learning strengthens cortical representations of motor sequences. Elife 2:e00801. doi: 10.7554/eLife.00801.

Willett FR, Deo DR, Avansino DT, Rezaii P, Hochberg LR, Henderson JM, Shenoy KV (2020) Hand knob area of premotor cortex represents the whole body in a compositional way. Cell 181:396–409.e26. doi: 10.1016/j.cell.2020.02.043.

Yarosh CA, Hoffman DS, Strick PL (2004) Deficits in movements of the wrist ipsilateral to a stroke in hemiparetic subjects. J Neurophysiol 92:3276–3285. doi: 10.1152/jn.00549.2004.

Yokoi A, Hirashima M, Nozaki D (2011) Gain field encoding of the kinematics of both arms in the internal model enables flexible bimanual action. J Neurosci 31:17058–17068. doi: 10.1523/JNEUROSCI.2982-11.2011.

Yokoi A, Hirashima M, Nozaki D (2014) Lateralized sensitivity of motor memories to the kinematics of the opposite arm reveals functional specialization during bimanual actions. J Neurosci 34:9141–9151. doi: 10.1523/JNEUROSCI.2694-13.2014.

Zhang CY, Aflalo T, Revechkis B, Rosario ER, Ouellette D, Pouratian N, Andersen RA (2017) Partially mixed selectivity in human posterior parietal association cortex. Neuron 95:697–708.e4. doi: 10.1016/j.neuron.2017.06.040.

Ziemann U, Hallett M (2001) Hemispheric asymmetry of ipsilateral motor cortex activation during unimanual motor tasks: further evidence for motor dominance. Clin Neurophysiol 112:107–113. doi: 10.1016/S1388-2457(00)00502-2.

