## supplementary materials for "Encoding of bimanual movement directions across the human sensorimotor system"

### Unimanual representational structure

To build our model components, we needed to establish the representational geometry for the six movement directions in the unimanual conditions. For contralateral and ipsilateral movements, the estimated second-moment matrix (using Eq. 2) was highly similar (Fig. S1a). The cosine similarity between the averaged contra and ipsilateral second-moment matrices was 0.82 ± 0.04 SEM across the sensorimotor regions. Therefore, for our modeling analysis, we averaged the unimanual second-moment matrix across the regions and contralateral/ipsilateral movements.

The representational geometry also provided some interesting insight into the encoding of wrist movement directions. When visualizing the movement pattern similarity separately for contra and ipsilateral movement (Fig. S1B,C), we found that movements in opposite directions were highly similar. We believe this mainly reflects the fact that activity was averaged across the out and back movements for each target. Additionally, we found that movements with wrist adduction (2,3) and wrist abduction (5,6) were overall more similar to each other.

When visualizing ipsi- and contralateral movement representations together in a single MDS plot (Fig. S1D), the correspondence of mirror-symmetric wrist movements also becomes visible. The shift in dimension 1 reflects the large mean activity difference between contralateral and ipsilateral movements. Despite this shift, however, the encoding of contra- and ipsilateral movement are aligned, providing a complementary representation of the strong intrinsic correlation found when considering the full-dimensional pattern (Fig. 4).

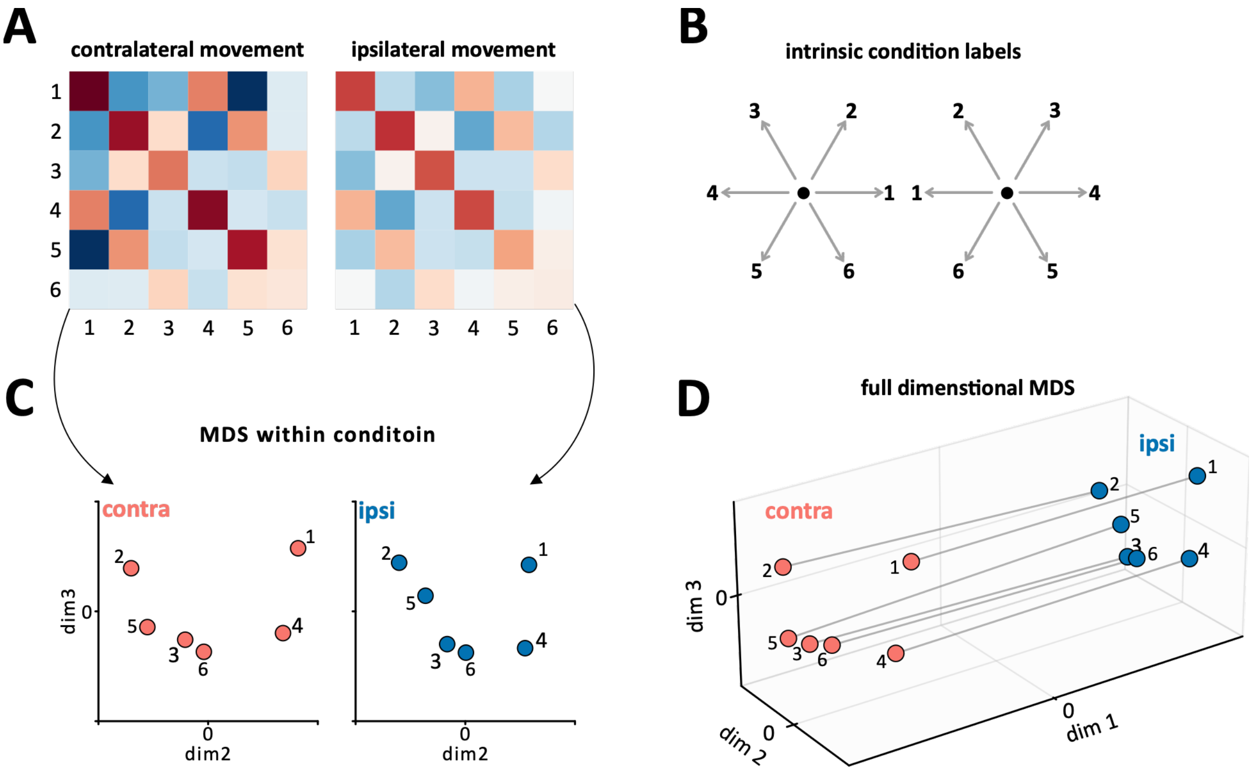


*Supplementary Figure 1:* ***Unimanual Structure***. ***A)*** *Double-centered cross-validated second-moment matrix of the contralateral and ipsilateral activity patterns averaged across ROIs, participants, hemispheres.* ***B)*** *The rows and column numbers in A correspond to the intrinsic numbering of the directions.* ***C)*** *Multidimensional Scaling (MDS) plot showing the relationship between the six directions within contra- (red) and ipsilateral (blue) activity patterns.* ***D)*** *The three-dimensional MDS view that captures the relationship between contra and ipsi activity patterns. dim2 and dim3 are the same directions as in C.*

### Hemispheric asymmetries in mean activity

Here we characterize the hemispheric differences in PSC (mean activity) within unimanual contralateral, ipsilateral and bimanual movement types across in all ROIs. All participants of this study were right-handed. For each movement type a 2-way repeated measures ANOVA tested for main effects of hemisphere side and ROI as well as their interactions. Finally, post-hoc two-sided paired t-tests were performed within each ROI.

*Contralateral movements.* A 2-way repeated measure ANOVA revealed significant main effect of ROI ($F_{6,66}$ = 21.52, p = 7.80e-14). The main effect of hemisphere was not significant ($F_{1,11}$ = 0.95, p = 0.35), however, there was a significant ROI-hemisphere interaction ($F_{6,66}$ = 3.17, p = 8.56e-3) suggesting that some ROIs might activate differently between the two hemispheres. Post-hoc t-tests revealed statistical significance only in SMA ($t_{11}$ = 4.64, $p_{fdr}$ = 5.03e-3), in which the right hemisphere was slightly more active. Overall, we did not find a statistically significant hemispheric asymmetry during contralateral movements in most ROIs except for SMA.

*Ipsilateral* movements. A 2-way repeated measure ANOVA revealed significant main effects of ROI ($F_{6,66}$ = 14.36, p = 2.15e-10), hemisphere ($F_{1,11}$ = 63.35, p = 6.85e-6), and a significant interaction ($F_{6,66}$ = 8.43, p = 8.56e-7). Post-hoc t-tests suggest that in most ROIs the left hemisphere was significantly more active than right hemisphere ($t_{11}$ > 4.01, $p_{fdr}$ < 2.82e-3, $p_{fdr}$ = 5.03e-3), except for PMv which did not reach statistical significance ($t_{11}$ = 1.89, $p_{uncorrected}$ = 0.085). SMA on the other hand was more active in the right hemisphere ($t_{11}$ = 2.32, $p_{uncorrected}$ = 0.041, $p_{fdr}$ = 0.048). Thus, the two hemispheres show asymmetries during ipsilateral movements.

*Bimanual* movements. A 2-way repeated measure ANOVA revealed significant main effects of ROI ($F_{6,66}$ = 11.08, p = 1.65e-8), hemisphere ($F_{1,11}$ = 9.10, p = 0.012), and a significant interaction ($F_{6,66}$ = 3.52, p = 4.41e-3). Post-hoc t-tests suggested significant differences in SMA ($t_{11}$ = 5.99, $p_{uncorrected}$ = 9.10e-5, $p_{fdr}$ = 6.37e-4) and the superior parietal ROIs ($t_{11}$ > 2.85, $p_{uncorrected}$ < 0.016, $p_{fdr}$ < 0.036), in which the right hemisphere was more active.

### Hemispheric asymmetries in encoding

We performed a similar comparison on the average cross-validated Euclidean distance (encoding) estimated within the contralateral, ipsilateral, and bimanual movements. In contrast to mean activity, we did not find any statistically significant differences between the hemispheres. Since our experiment was heavily focused on the bimanual activity patterns, the number of trials sampled from unimanual contralateral and ipsilateral movements might not be large enough to depict the hemispheric asymmetries that might exist in encoding.
